# Multiomic Insights into Human Health: Gut Microbiomes of Hunter-Gatherer, Agropastoral, and Western Urban Populations

**DOI:** 10.1101/2024.09.03.611095

**Authors:** Harinder Singh, Rosana Wiscovitch-Russo, Claire Kuelbs, Josh Espinoza, Amanda E. Appel, Ruth J. Lyons, Sanjay Vashee, Hagen E.A. Förtsch, Jerome E. Foster, Dan Ramdath, Vanessa M. Hayes, Karen E. Nelson, Norberto Gonzalez-Juarbe

## Abstract

Societies with exposure to preindustrial diets exhibit improved markers of health. Our study used a comprehensive multi-omic approach to reveal that the gut microbiome of the Ju/’hoansi hunter-gatherers, one of the most remote KhoeSan groups, exhibit a higher diversity and richness, with an abundance of microbial species lost in the western population. The Ju/’hoansi microbiome showed enhanced global transcription and enrichment of complex carbohydrate metabolic and energy generation pathways. The Ju/’hoansi also show high abundance of short-chain fatty acids that are associated with health and optimal immune function. In contrast, these pathways and their respective species were found in low abundance or completely absent in Western populations. Amino acid and fatty acid metabolism pathways were observed prevalent in the Western population, associated with biomarkers of chronic inflammation. Our study provides the first in-depth multi-omic characterization of the Ju/’hoansi microbiome, revealing uncharacterized species and functional pathways that are associated with health.

## Introduction

For the past 10,000 years, the human way of life and diet have drastically transformed due to the development of agriculture and even more after the Industrial Revolution^1^. The rapid shift from a Paleolithic to a Western diet has disrupted highly efficient metabolic pathways that evolved over millions of years of human and host-associated microbial dietary adaptation, leading to the development of Western diseases (WD)^2^. WDs are defined as non-infectious chronic degenerative diseases that are rare or absent in agropastoral (AP) and hunter-gatherer (HG) populations^3^. They are the leading causes of morbidity and mortality in Western populations^3^. WD includes chronic inflammation, obesity, type 2 diabetes (T2D), cardiovascular diseases (CVD), and cancer^4^. In contrast, infectious diseases are the primary cause of death in AP and HG populations^3^. In addition, several studies of HG populations have observed low circulating insulin levels and optimal insulin sensitivity if they maintain their traditional diet^5^. These observations underline the importance of considering the evolutionary history of human dietary patterns in the context of modern dietary practices to promote optimal health outcomes^2^.

In the past decade, our understanding of the relationship between human health and the gut microbiome has expanded significantly. It is now well-established that the gut microbiome is crucial for overall health, influencing conditions such as nutrition deficiencies, obesity, chronic inflammation, diabetes, and even cancer^6^. It regulates energy harvest, signaling molecule release, and plays an invaluable role in maintaining the intestinal barrier and immune cell repertoire in the mucosa^7–9^. Understanding the gut microbiome’s complexities, its relationship with human health, and the factors that shape its composition is critical for the development of effective therapeutic strategies that can mitigate chronic diseases. Due to recent significant studies from the National Institutes of Health-Human Microbiome Project and the European Metagenomics of the Human Intestinal Tract, our understanding of the genes, function, and complexity within the composition of the human gut bacteria has become more complete^10,11^. Notably, the microbiome’s diversity plays a pivotal role in facilitating the production of nutrients from substrates that the host cannot metabolize^12^. Differences in gut microbiota between Western urban populations, HG, and AP are driven by dietary variations^13^, with seasonal dietary dependencies causing significant microbiome shifts in groups such as the Hazda HG of Tanzania^14,15^. Western populations have less diverse gut microbiota, which lacks organisms that promote health and metabolically efficient digestion. In contrast, HG and AP have more diverse and stable gut microbiota due to their high-fiber diets and low-fat and sugar consumption^15,16^. These differences have significant health implications, since alterations in gut microbiota composition and diversity are linked to chronic diseases such as obesity, type 2 diabetes, and inflammatory bowel disease^14,16–18^.

The modern human originated in Africa over 200 thousand years ago and primarily relied on a hunter-gatherer’s lifestyle when the everyday diet consisted mainly of hunted wild animals, fish, and unfarmed plant-based foods^19,20^. Of significant importance, a recent report has shown that anatomically modern humans originated from the Kalahari Desert region in Namibia^21^. The Ju/’hoansi community in the Kalahari Desert represents a unique and remote segment within the ethnolinguistic KhoeSan framework. Furthermore, the Ju/’hoansi are among the ‘oldest’ extant (or earliest diverged) contemporary human populations, pre-dating the Baka Pygmy divergence^22^. One of the oldest hunter-gatherer cultures that reside in the Kalahari Desert, the Ju/’hoansi provide a living example of a foraging and hunting lifestyle, with a plant-centric diet rich in ∼105 edible plants consisting primarily of nuts, fruits, roots, gums, and leafy greens with meat being scarce^23,24^. However, due to the seasonal scarcity of natural resources, the Ju/’hoansi now depend on a mix of traditional hunter-gatherer methods and market-based strategies. Of note, a recent study has provided a low-depth V3-V4 16S rRNA and ITS1 sequencing for bacterial and fungal identification, respectively, from the Ju/’hoansi living at Nyae Nyae community^25^. While lacking species-level identification, this study documented the microbiota of Ju/’hoansi and established the unique role of diet and culture in core microbial composition compared to urban populations. Here, we discern the composition and function of the microbiome of geographically related HGs and APs and subsequently compare them to a Western diet cohort of genetically similar descendants from the West Indies.

For this research, we collected fecal samples from the Ju/’hoansi population residing in the Kalahari Desert of Northern Namibia, the Bantu AP group in the nearby vicinity, and a cohort comprising both healthy and diabetic/obese (defined as any participant with BMI ≥30 kg.m^2^, or with an HbA1c value of ≥6.5%) individuals from Trinidad (West Indies) and then conducted extensive metagenomic, metatranscriptomic, and metabolomics sequencing to analyze their gut microbiome profiles. Our metatranscriptomics approach offers one of, if not, the most deeply sequenced human gut microbiomes to date (∼150 million paired reads on average). Moreover, we obtained a significant amount of high-quality metagenomic and metabolomic data, including 97 GB of metagenomic assemblies, 9,137 high-quality assembled genomes, 916 global metabolites, and 8 short-chain fatty acids abundance data. This data allowed us to identify microbes, microbial genes/pathways, and metabolites that are highly abundant or absent in the Ju/’hoansi population compared to the Western urban population. The data also revealed the presence/absence of microbes, microbial genes/pathways, and metabolites associated with WD that are not associated with the Ju/’hoansi population. All data we generated is freely available for future study by other scientists, including (1) metagenomic sequencing reads, (2) high-quality metagenome-assembled genomes (MAGs), (3) metatranscriptomics sequencing reads, (4) metabolomics data, and (5) organized metadata.

## Results

### A multi-omics approach to study complex microbiomes

In this study, a total of 219 fecal samples were collected for analysis from 55 Ju/’hoansi HG, 16 Bantu AP, 90 healthy and 58 obese/diabetic (BMI≥30kg.m^2^ and HbA1c value of ≥6.5%) people of African descent from urban Trinidad, respectively (**Figure 1**). To perform ultra-high-depth metatranscriptomics (MetaT) and a high-depth sequencing approach for metagenomics (MetaG), we used the Illumina NovaSeq 6000 system. After the exclusion of human data, MetaT yielded approximately 30.4 billion microbial read pairs. Similarly, for MetaG, the removal of human data resulted in 10.5 billion microbial reads. MetaSPAdes was then used to generate 219 assemblies totaling 96.6 GB of assembled data. Using a combinatorial bioinformatic approach^26^ (**Figure 1**), we identified 9,010 bacterial MAG, 127 archaeal genomes with 726 genomes not classified at the species level, 1025 viral genomes, and 2 eukaryotic genomes. After performing whole-genome average nucleotide identity-based clustering using FastANI, the final clusters delineated 1,184 bacterial species-level clusters (SLC), 7 archaeal SLC, 650 viral (bacteriophage) SLC, and 2 eukaryotic SLC (**Table S1**). The bar plot in Figure 1 displays the counts of genomes at the phylum level. Blue-colored MAGs represent organisms classified at the species level, while those not classified at the species level are depicted in orange using the GTDB-Tk^27^. The MetaT and Taxonomical analysis assessment involved utilizing the VEBA pipeline, a computational pipeline designed for metagenomic analysis ^26^. Gene Set Enrichment Analysis (GSEA) was also employed to identify enzymes, biological pathways, or processes overrepresented in the analyzed gene sets^28^. The set of 916 global metabolites offers a meticulously curated analysis of metabolon pathways, encompassing both super-pathways and sub-pathways. Super-pathways are composite biochemical pathways that elucidate the biosynthesis or metabolism of interconnected compounds, while subp-athways represent a segment of biochemical pathways. In summary, while the microbiome analysis of the combined samples showed mapping to a vast number of microbes present in the Genome Taxonomy Database^29^, many of the recovered genomes were absent in such databases, detailing the novelty of high-depth-sequencing approaches for the discovery of unique microbial groups (**Figure 1, and Table S1**).

**Figure 1:**
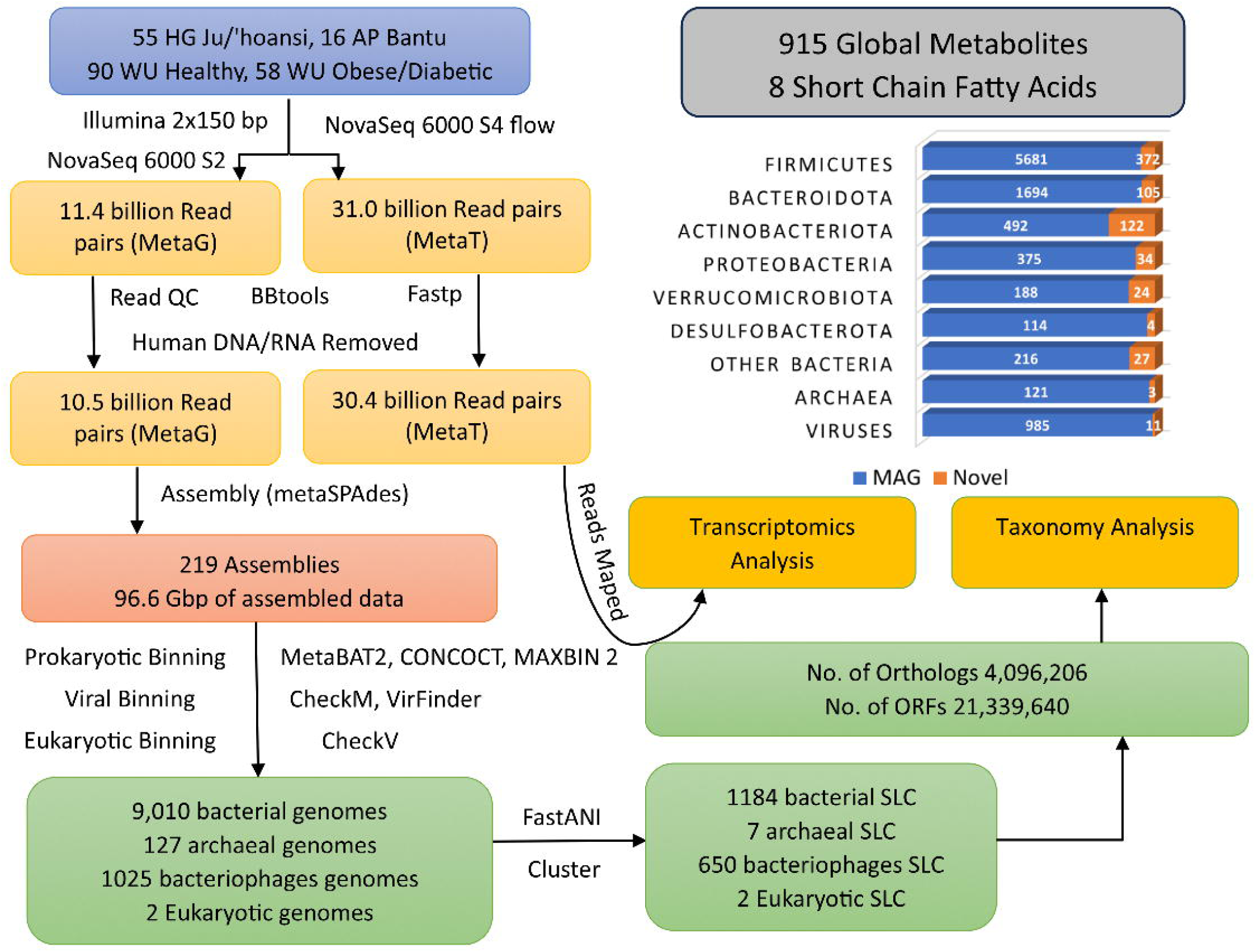
Resource for human gut microbiome evolution. The computational workflow used to generate high-quality metagenome-assembled genomes (MAGs) from Illumina metagenomic (MetaG) and metatranscriptomics (MetaT) sequencing. Blue MAGs represent classified organisms, while unclassified ones appear in orange using GTDB-Tk.

### The gut microbiome of the Ju/’hoansi people

First, we assessed the microbial abundance of the Ju/’hoansi gut microbiome. The circular representations of the taxonomic and phylogenetic tree of 1,184 bacterial species clusters present among all four groups are shown in **Figure 2A**. The size of clade markers inside the circular plot represents the relative abundance of the bacterial clusters in the Ju/’hoansi gut microbiome. The Ju/’hoansi gut microbiome is mainly composed of bacteria from the Phyla of Firmicutes (47.5%), Bacteroidota (39.8%), Proteobacteria (4.9%), Spirochaetota (2.8%), and Verrucomicrobiota (1.8%). Although Firmicutes is the most abundant and diverse phyla, genera from Bacteroidota (*Cryptobacteroides, Prevotella, Parabacteroides, Bacteroides*), Proteobacteria (*Succinivibrio*) and Spirochaetota (*Treponema_D)* are the most abundant genera. Among the most abundant classified SLC are *Succinivibrio sp000431835* (3.1%)*, Cryptobacteroides sp000433355* (2.8), *Prevotella copri A* (1.9%), *Prevotella sp900313215* (1.8%), *Parabacteroides sp900549585* (1.7%), *Prevotella sp900546535* (1.3%), *Prevotella sp900556795* (1.8%), *Cryptobacteroides sp000432515* (1%), *Treponema D succinifaciens* (1%) and *Bacteroides uniformis* (1%) (**Figure 2A**). Additional prokaryotes found in the Ju/’hoansi gut microbiome were Archaea, comprising less than 1% of the microbiome. The most abundant Archaea species in the Ju/’hoansi were *Methanomassillicoccales intestinalis*, *Methanobrevibacter smithii*, *Methanosphaera stadtmanae, Methanomethylophilus alvus* (**Figure 2B**). In the case of eukaryotes, only two metagenome-assembled genomes (MAGs) belonging to the species subtype 4 of Blastocystis sp. and the genus Blastocystis were identified. In the case of DNA/RNA viruses, we observed a major abundance of bacteriophages from the Caudovirales, Microviridae, and Martellivirales (**Figure 2C**).

**Figure 2:**
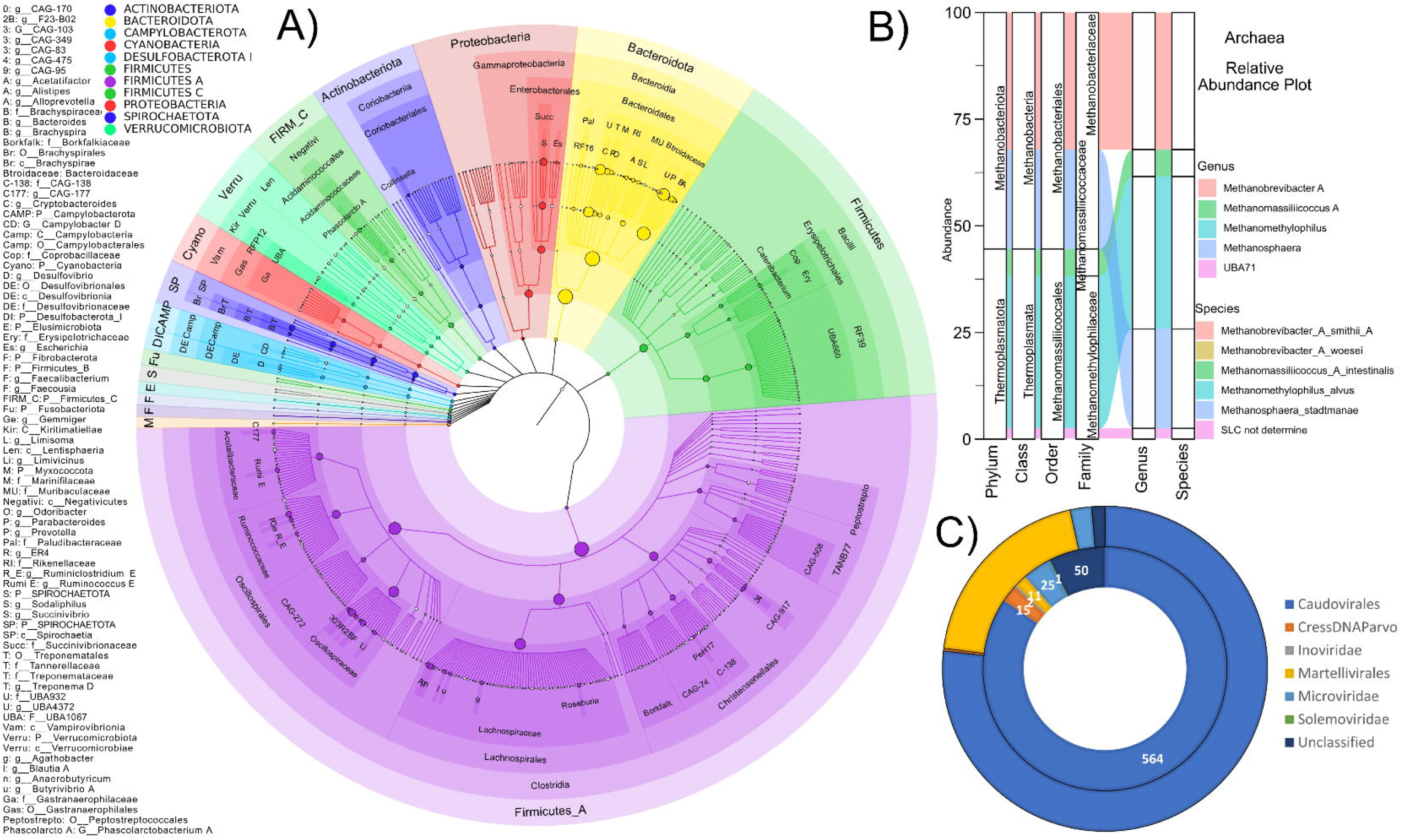
Gut microbiome diversity and abundance profile of Ju/’Hoansi peoples. **A**) The bacterial SLC relative abundance as circular representations of taxonomic and phylogenetic trees. Each color represents a different phylum, and the circle size represents the abundance of the respective taxonomic level. **B**) Relative abundance distribution across different taxonomic levels of Archaea SLC. **C**) The visualization illustrates the frequency (inner circle) and relative abundance of different viral/phage SLC (outer circle).

### Microbiome of Ju/’hoansi people is distinct from Western urban African descendants

When comparing the microbiome of the rural population (Ju/’hoansi and Bantu) and the Western urban (WU, healthy and obese/diabetic), we observed 620 significant (adjusted p-value <= 0.05) differentially abundant SLC between the groups. Among these 620 SLC, 269 are more abundant in the rural population, with 132 SLC having an effect size more than 1.0, and 351 were more abundant in the WU population, with 230 SLC with an effect size greater than -1.0 (**Figure 3A-B, Table S1**). The Ju/’hoansi and Bantu showed higher alpha diversity than the WU healthy group, followed by the obese/diabetic group (**Figure 3C**). The principal component axis (PCA) plot shows a separate clustering of Ju/’hoansi, Bantu, WU healthy and obese/diabetic cohorts, with Bantu showing a 95% confidence interval ellipse overlapping healthy and obese/diabetic cohorts (**Figure 3D**). At the phylum level, Spirochaetota and Campylobacterota were the most abundant in the Ju/’hoansi (1.53, 0.96 effect size) population, followed by Bantu (0.25, 0.18), WU healthy (-0.55, -0.34) and WU obese/diabetic (-0.34, -0.35). The least abundant phyla in Ju/’hoansi were Actinobacteriota and Firmicutes A (-1.64, -1.18), with increasing abundance in Bantu (-1.24, -0.71), WU obese/diabetic (0.47, 0.45) and WU healthy (0.94, 0.67), respectively. Overall, six phyla were more abundant (1.53 to 0.43), and thirteen phyla were less abundant in the Ju/’hoansi as a whole (-0.03 to -1.64) when compared to the other populations (**Figure 3B, 3E, Table S2**).

**Figure 3:**
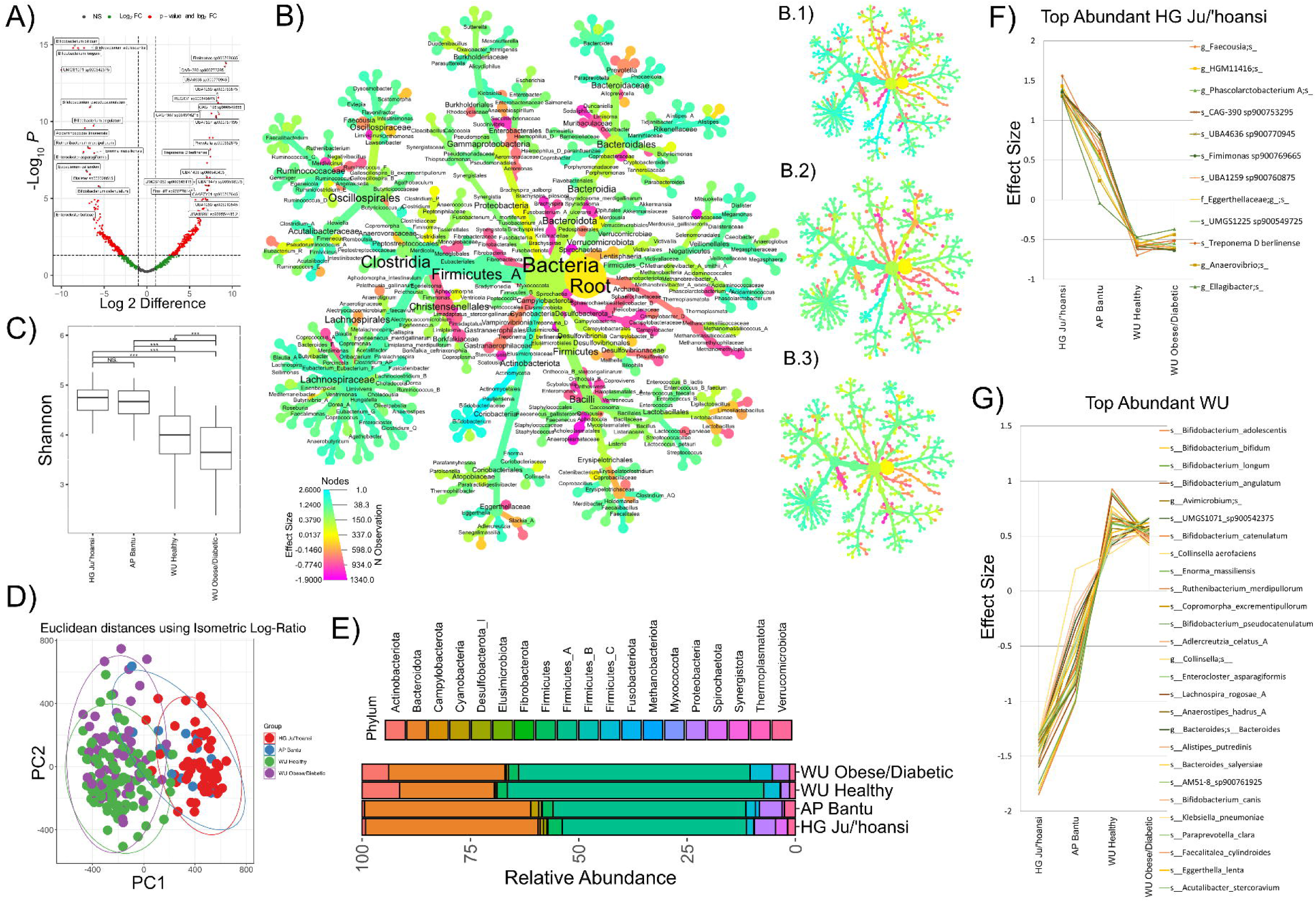
Microbiome comparison of Ju/’hoansi, Agropastoral, Urban Healthy and Urban obese/diabetic population. **A)** Volcano plot showing the differentially abundant species between rural and WU using ALDEx2. **B)** Multiple heat trees showing significant differentially abundant kingdom, phylum, class, order, family, genus, and species in the Ju/’hoansi population (Benjamini-Hochberg corrected p-value using Wilcoxon test < 0.05) with labels up to genus are displayed. The AP Bantu population is displayed in **(B.1)**, WU healthy in **(B.2)**, and WU obese/diabetic in **(B.3). C)** The Alpha Diversity plot uses the Shannon diversity index to compare the diversity of four cohorts. **D)** PCA plot generated using isometric log-ratio data based Euclidean distance plot between the four cohorts. **E)** The phylum-level relative abundance composition bar plot shows the distribution of various phylum in four cohorts. **F)** The topmost species with decreasing abundance from Ju/’hoansi to AP Bantu to WU cohort. **G)** The topmost species with increasing abundance from Ju/’hoansi to AP Bantu to WU cohort.

When looking at the differences in the topmost abundant bacterial species, an increase in *UMGS1225 sp900549725, CAG-390 sp900753295, UBA4636 sp900770945, Fimimonas sp900769665, UBA1259 sp900760875, Treponema D berlinense* and several unclassified species from the genera *Faecousia, HGM11416, Phascolarctobacterium A, Eggerthellaceae, Anaerovibrio, Ellagibacter* (effect size from 1.56 to 1.3) was observed in Ju/’hoansi (**Figure 3A, Table S3**). Among other abundant genera with effect size >1.0 are: *Anaerovibrio, Aphodocola, Cryptobacteroides, ER4, Faecousia, Fimimonas, Gemmiger, Limivicinus, Onthousia, Onthovivens, Phascolarctobacterium A, Prevotella, Scatousia, Stercorousia,* and several unclassified/uncultured genus/species belong to class Acholeplasmatales, Acidaminococcales, Bacteroidales, Christensenellales, Coriobacteriales, Elusimicrobiales, Erysipelotrichales, Gastranaerophilales, c_Bacilli;o_ML615J-28, Monoglobales, Oscillospirales, Peptostreptococcales, c_Alphaproteobacteria;o_RF32, c_Bacilli;o_RF39, c_Bacilli; RFN20, c_Kiritimatiellae;o_RFP12, Selenomonadales, c_Clostridia;o_TANB77. The majority of prevalent species in the Ju/’hoansi community remain either unclassified or uncultured. (**Figure 3B, Table S3**). Of interest, microbial groups with the highest abundance in the Ju/’hoansi sequentially decreased in the Bantu and the WU healthy and obese/diabetic populations, respectively (**Figure 3A, 3F, Table S3**).

Common probiotic species such as *Bifidobacterium catenulatum, Bifidobacterium angulatum, Bifidobacterium longum, Bifidobacterium bifidum, Bifidobacterium adolescentis, Bifidobacterium pseudocatenulatum* were significantly reduced and/or absent in the rural population when compared to the WU population (**Figure 3A, 3G, Table S3**). Other species with effect size greater than -1.3 in the Ju/’hoansi were *Eggerthella lenta, Faecalitalea cylindroides, Klebsiella pneumoniae, Paraprevotella clara, Alistipes putredinis, Bacteroides salyersiae, Anaerostipes hadrus A, Lachnospira rogosae A, Ruthenibacterium merdipullorum, Copromorpha excrementipullorum, Enorma massiliensis* and species from *Adlercreutzia, Alistipes, Anaerostipes, Avimicrobium, Bacteroides, Bifidobacterium, Collinsella, Copromorpha, Eggerthella, Enorma, Enterocloster, Faecalitalea, Lachnospira, Paraprevotella, Ruthenibacterium, f_Acutalibacteraceae;g_UMGS1071, f_Lachnospiraceae;g_AM51-8* genera (**Figure 3A, 3G, Table S3**). This data conclusively demonstrates the gut microbiome profile of the Ju/’hoansi is distinct from the AP and WU populations, highlighting the significance of this unique microbiome.

### Microbial gene expression in the Ju/’hoansi compared to the Bantu and WU populations

We also characterized the microbial gene expression of the gut microbiome by utilizing MetaT. Among the 620 differentially abundant SLCs, 72,055 orthologs were present in 70% of either the rural or the WU population. Using these 72,055 orthologs, we found 48,114 orthologs differentially abundant between the rural and WU groups (adjusted p-value <= 0.05). From these 48,114 orthologs, we calculated the NES (normalized enrichment score) for various KEGG pathways, modules, ortholog groups, and BRITE hierarchies. There were 141 KEGG orthologs with significant NES scores (adjusted P value < 0.05), 50 KEGG orthologs with NES below -1, and 91 with NES above +1 (**Figure 4A, Table S4)**. The KEGG orthologs with the highest expression in the rural population (Ju/’hoansi and Bantu) as compared to the WU population (healthy and obese/diabetic) included pyruvate-ferredoxin, elongation factor G/Tu, membrane protein, pyruvate orthophosphate dikinase, DNA-directed RNA polymerase, phosphoenolpyruvate carboxykinase, peptide transport system permease, iron (III) transport system substrate-binding protein, DNA gyrase subunit A, DNA-directed RNA polymerase subunit, preprotein translocase subunit SecY, glyceraldehyde 3-phosphate dehydrogenase, topoisomerase IV subunit A, diphosphate reductase, aldehyde oxidoreductase, H+/Na+-transporting ATPase subunit A/B, oligoendopeptidase F, glucose-1-phosphate adenylyltransferase, CO dehydrogenase maturation factor, iron (III) transport system permease protein, maltose regulon positive regulatory protein, phosphoenolpyruvate carboxykinase, NAD+ oxidoreductase, chaperonin GroEL, methyl-accepting chemotaxis protein, fructose-bisphosphate aldolase, starch synthase, and both small and large ribosomal proteins (**Figure 4A, Table S4**).

**Figure 4:**
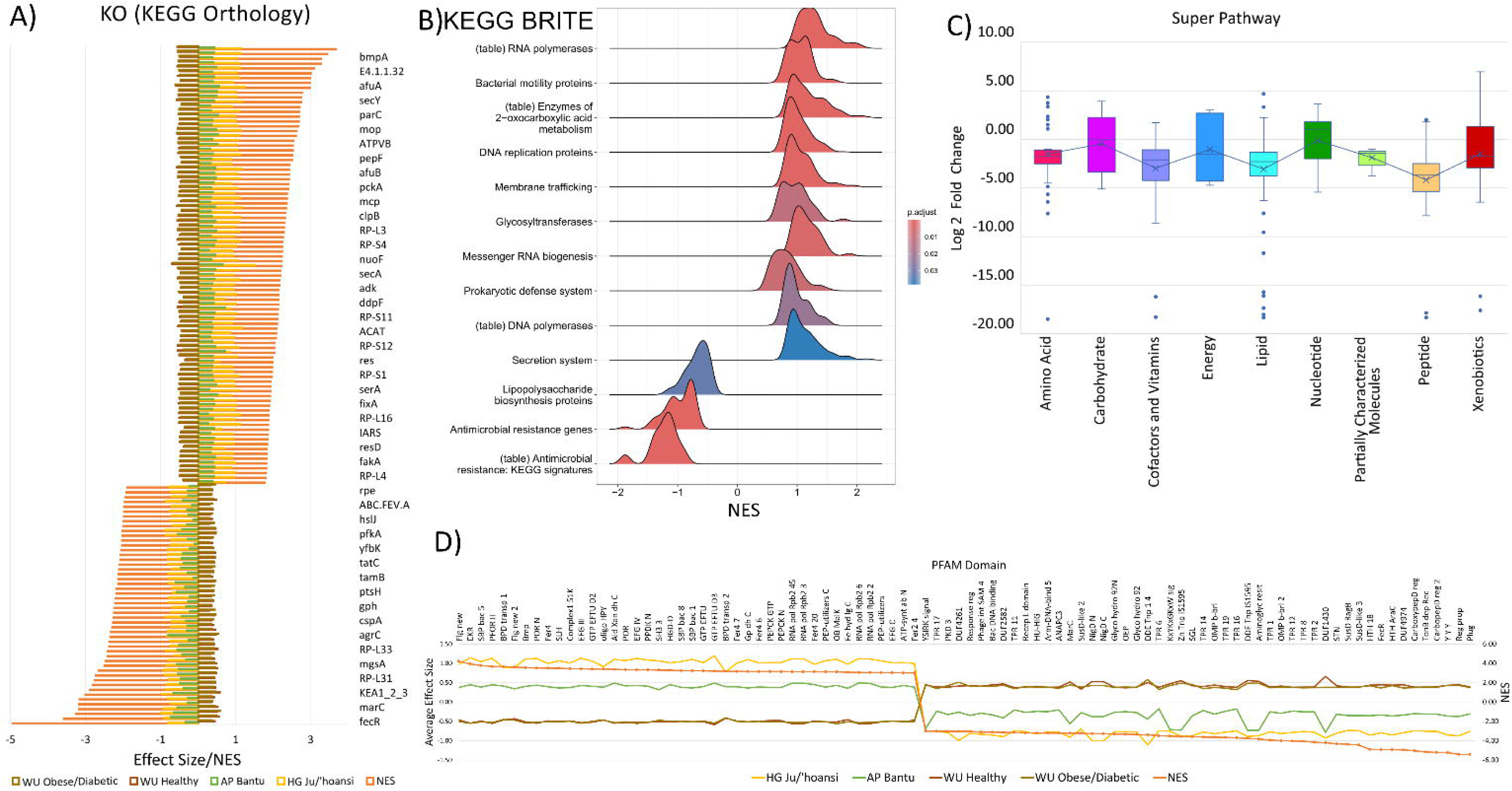
Metatranscriptomics comparison overview: **A)** NES (normalized enrichment score) for the top KEGG orthologs with corresponding effect size across four cohorts plotted as a bar plot. **B)** NES score for KEGG BRITE hierarchies as x-axis and adjusted p. Value as color code. **C)** Super pathway box plot displaying all metabolites’ log 2-fold change (> 1.0). **D)** The Pfam domain with the highest NES scores and the effect size of four cohorts is displayed as a line graph.

At the KEGG BRITE hierarchy level, gene activity related to RNA polymerases, bacterial motility proteins, enzymes of 2-oxocarboxylic acid metabolism, DNA replication proteins, membrane trafficking, glycosyltransferases, messenger RNA biogenesis, prokaryotic defense system, DNA polymerases, and secretion system was more active in the rural populations. (**Figure 4B**). Of the 547 Pfam domains, 293 have NES scores higher than +1, whereas 254 Pfam domains showcase NES scores lower than -1 (**Table S5**). The Pfam domains more expressed in the rural population include multiple pyruvate ferredoxin/flavodoxin oxidoreductase domains, extracellular solute-binding proteins, inner membrane component, ABC transporter, multiple iron/4Fe-4S binding domain, S-layer homology domain, NADH dehydrogenase, Elongation Factor G/Tu, Oligopeptide transporter, Aldehyde oxidase and xanthine dehydrogenase, Bacterial SH3 domain, CoA dehydratase, amino acid transport system, Glyceraldehyde 3-phosphate dehydrogenase, various RNA polymerase domains, PEP-utilising enzyme, MalK OB fold domain, and ATP synthase alpha/beta family. The global metabolites profile comparison shows that metabolites from carbohydrate and nucleotide superpathways exhibit comparable abundance levels in both rural and urban populations. (**Figure 4D**).

The KEGG orthologs with the highest expression in the WU population included Fe^2+^ transmembrane sensor, transposase, macrolide efflux protein, multiple antibiotic resistance proteins, iron complex outer membrane receptor, periplasmic protein TonB, K+:H+ antiporter, RNA polymerase sigma-70 factor, acyl carrier protein, phosphoglycerate mutase, ferritin, methylglyoxal synthase, LemA protein, REP-associated tyrosine transposase, beta-fructofuranosidase, sensor histidine kinase AgrC, biopolymer transport protein ExbD, regulator for asnA, asnC and gidA, cold shock protein, response regulator YesN, phosphoglycolate phosphatase, transcription termination/antitermination protein NusG, starch synthase, and large subunit ribosomal proteins (**Figure 4A, Table S4**). At the KEGG BRITE hierarchy level, genes involved in lipopolysaccharide biosynthesis proteins and antimicrobial resistance genes were more active in the WU population (**Figure 4B**). Pfam domains more expressed in the WU populations include: Two component regulator propeller, TonB-dependent Receptor, Carboxypeptidase regulatory-like, Bacterial regulatory helix-turn-helix proteins, FecR protein, Starch-binding, SusD family, multiple Tetratricopeptide repeat domain, many Outer membrane proteins domain, Aminoglycoside-2’’-adenylyltransferase, ISXO2, Gluconolactonase, Transposase zinc-ribbon, Glycosyl hydrolase, NigD domain, Anaphase-promoting complex MarC family, Arm DNA-binding domain, HU domain, Winged helix-turn-helix, Phage integrase, and many signal peptides (**Figure 4C, Table S5**). The global metabolites profile comparison shows that metabolites in amino acid, lipid, cofactors and vitamins, partially characterized molecules, and peptides superpathway were more active in the WU population with log 2-fold changes ranging from -1.01 to -17.60. The average log 2-fold change of Energy and Xenobiotics superpathways metabolites demonstrates a tendency toward the WU population (**Figure 4D**).

### The highly active and defensive Ju/’hoansi gut microbiome lacks the presence of major antimicrobial resistance genes and several typical food metabolites

The Ju/’hoansi gut microbiome displayed significant metabolic activity and a heightened resilience to stress-inducing environments when contrasted with the Bantu and WU population gut microbiome, showing an effect size twice for respective microbial active genes (**Figure 5B**). Such active pathways and enzymes include DNA-directed RNA polymerase, Bacterial motility proteins, Trehalose biosynthesis, Quorum sensing, ABC transporters, Flagellar assembly, Membrane trafficking, RNA degradation, Mismatch repair, DNA replication, mRNA biogenesis, Bacterial secretion system, Protein export and Secretion system (**Figure 5B**). The Ju/’hoansi group has exhibited significantly higher Trehalose biosynthesis pathway activity than the WU population. Trehalose, a stable disaccharide molecule, is a source of energy and carbon metabolism when responding to abiotic stresses and has been shown to regulate blood glucose levels^30–34^. In addition, the rural population exhibited elevated RNA-based microbial activity, as evidenced by observed alterations in nucleotide metabolite levels compared to the WU population (**Figure 5C**).

**Figure 5.**
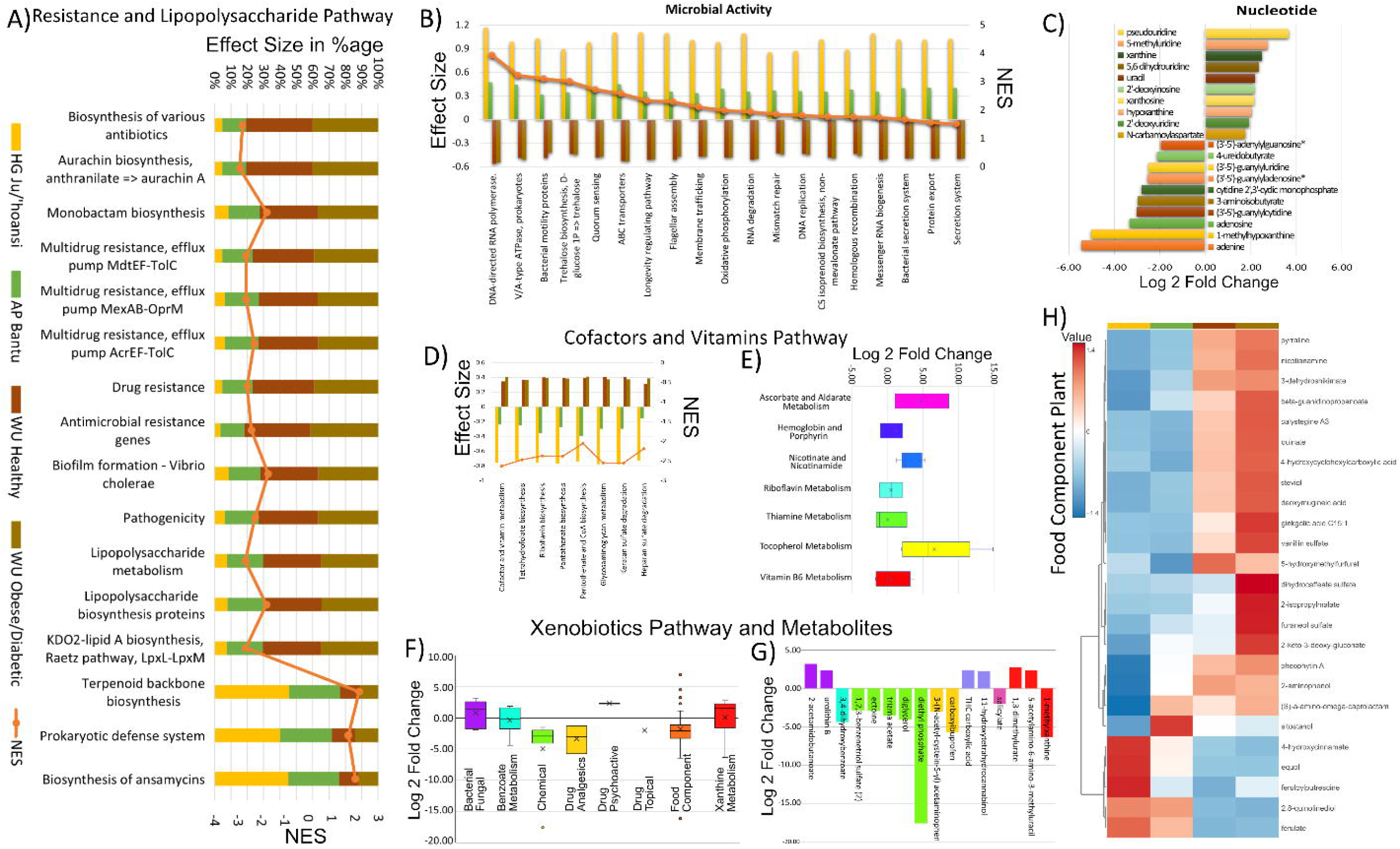
Microbial resistance, activity, and food metabolism. **A)** Antibiotic resistance and prokaryotic defense system pathways. The four-bar color ratio represents the proportion of the total effect size, and the orange line denotes the NES score for the respective pathway. **B)** Various microbial activity pathways, each bar representing the effect size and the orange line denoting the NES score for the respective pathway. **C)** Nucleotide metabolites log 2-fold changes with negative values representing the log-fold change in the WU population. **D)** Microbial genes and **E)** metabolites involved in the cofactor and vitamin pathway based on metatranscriptomics and metabolomics analysis. **F)** The abundance of the Xenobiotic pathway and **G)** metabolites in the WU and rural populations. **G)** The Food Metabolites heatmap shows the abundance of significant food metabolites in four cohorts.

Among cofactors and vitamins subpathways, metabolites from ascorbate and aldarate metabolism, a crucial carbohydrate metabolic pathway that protects cells from oxidative damage^35^, nicotinate, and nicotinamide metabolism, which plays a central role in cellular energy metabolism, DNA damage repair, gene expression, and stress response^36^ were highly abundant (18.29 log fold change of NAD+) in the rural compared to WU population. Other metabolite subpathways, including hemoglobin and porphyrin metabolism, riboflavin metabolism, thiamine metabolism, and vitamin B6 metabolism, showed similar abundance in rural and WU populations (**Figure 5E**). The Ju/’hoansi have a high abundance of metabolites from the byproducts of the Bacterial Fungal subpathway, including 2-acetamidobutanoic acid, Urolithin B (improves muscle function)^37,38^, and glutamyl-meso-diaminopimelate, diaminopimelate which are involved in peptidoglycan synthesis^39^ (**Figure 5G**). The rural population surprisingly exhibits a high presence of two caffeine metabolites, namely 5-acetylamino-6-amino-3-methyluracil and 1,3-dimethyluric acid^40^, along with metabolites of cannabis such as THC carboxylic acid and 11-hydroxytetrahydrocannabinol^41^ (**Figure 5F, G)**. Among plant food metabolites, ferulate and 2,8-quinolinediol were also highly abundant in the rural groups. Three plant metabolites, equol, 4-hydroxycinnamate, and Feruloylputrescine, were abundant only in the Ju/’hoansi (**Figure 5G)**. They possess diverse bioactive properties, including antioxidant, antimicrobial, anticancer, anti-inflammatory, anti-diabetic, anti-melanogenic, and cardioprotective effects, and can inhibit aflatoxin production ^42–48^.

In the case of the WU population, a significant enrichment of various antimicrobial resistance (AMR) pathways was observed in the WU cohort. These include efflux pump MdtEF-TolC rhodamine 6G, which confers resistance against erythromycin, doxorubicin, and ethidium bromide^49–52^; the efflux pump MexAB-OprM, which confers resistance against carvacrol, quinolones, macrolides, novobiocin, chloramphenicol, tetracyclines, lincomycin, and β-lactam antibiotics^53,54^, and the efflux pump AcrEF-TolC, which confers resistance against fluoroquinolone, quinolone, ciprofloxacin^55–58^. The WU was also enriched in aurachin A, a type II polyketide that inhibits the respiratory chain in prokaryotes and eukaryotes, as well as Monobactam, which inhibits peptidoglycan synthesis process biosynthesis pathways (**Figure 5A**)^59^. The WU population exhibits elevated expression levels of various metabolic pathways, notably including cofactor and vitamin metabolism, tetrahydrofolate biosynthesis, riboflavin biosynthesis, pantothenate, CoA biosynthesis, and glycosaminoglycan metabolism, including heparan sulfate degradation. These pathways were found to be more prevalent in probiotic bacteria, which are in turn more abundant in the WU population, particularly within the healthy cohort (**Figure 5D**)^60^.

Metabolite subpathways, including hemoglobin and porphyrin metabolism, riboflavin metabolism, thiamine metabolism, and vitamin B6 metabolism, showed similar abundance in rural and WU populations (**Figure 5E**). Xenobiotic pathways including chemical, drug analgesics, food metabolites and several metabolites including some drugs were abundant in the WU population, including ectoine, acetaminophen, trizma acetate, diglycerol, 3,4-dihydroxybenzoate, carboxyibuprofen, 1-methylxanthine, and diethyl phosphate (**Figure 5F, G)**. Diethyl phosphate is a neurotoxin^61^, diglycerol is used as a fat restriction in the management of diabetics^62^, it is a food stabilizer, and triethanolamine is found in cosmetic products, herbicides, and algicide^63^. Ectoine helps survival under extreme osmotic stress^64^ (**Figure 5G**). The food metabolites prevalent in the WU population, especially in the obese/diabetic group, were advanced glycation end products (AGEs), which are biomarkers of aging, diabetes, diabetic ketoacidosis, and chronic kidney disease. These metabolites consist of pyrraline, nicotianamine, 3-dehydroshikimate, beta-guanidinopropanoate, calystegine A3, quinate, 4-hydroxycyclohexylcarboxylic acid, steviol, deoxymugineic acid, ginkgolic acid C15:1, vanillin sulfate, 5-hydroxymethylfurfural, dihydrocaffeate sulfate, 2-isopropylmalate, furaneol sulfate, 2-keto-3-deoxy-gluconate, pheophytin A, and 2-aminophenol. These metabolites are intermediates in amino acid biosynthesis pathways, aromatic bioproducts, and industrial chemicals. They are naturally present in foods such as potatoes, peppers, paprika, eggplants, strawberries, pineapples, tomatoes, buckwheat, nutritional supplements, and flavoring compounds used in foods, beverages, and pharmaceuticals. Some metabolites have antioxidant, potential allergenic properties, and cytotoxic and carcinogenic effects ^65–76^ (**Figure 5F, 5G**).

### Low Amino acid, Fatty acid, and lipid metabolism pathway expression in the Ju/’hoansi

The abundance and expression of amino acid pathways were significantly downregulated in rural populations (**Figure 6A-C**). Metabolites derived from the metabolism of leucine, isoleucine, and valine display heightened abundance within the rural population (**Figure 6B and Table S6)**. At the transcriptomics pathway expression level, branched-chain amino acids (BCAAs), including the leucine, isoleucine, and valine degradation pathway, were found more enriched in the Ju/’hoansi population, which supports the high abundance of leucine amino acids and peptides (**Figure 6C and Table S7**). The rural population exhibits abundance in the pathways of Phosphatidylcholine PC and Ceramides lipids (**Table S8)**. In rural areas, the most prevalent lipid was oleoyl-linoleoyl-glycerol, which was found in high abundance (**Table S8)**. Finally, a line plot delineates the prevalence of bacterial species, including novel ones, that have enhanced expression of Leucine, Valine, and Isoleucine pathways in the Ju/’hoansi and the Bantu populations and not the WU population (**Figure 6F)**.

**Figure 6.**
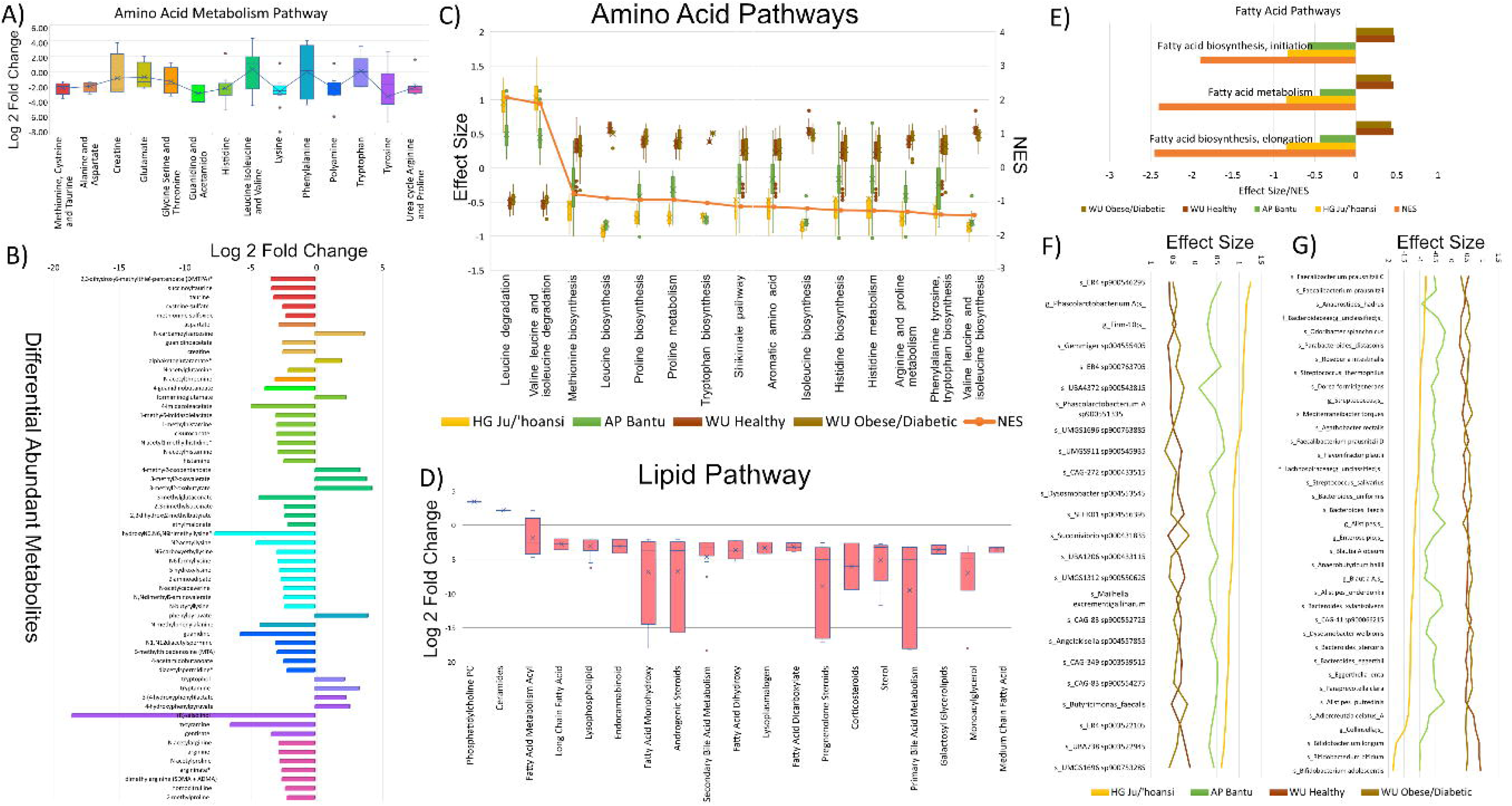
**A)** Box plot illustrating the log 2-fold changes for all amino acid metabolism pathway metabolites. **B)** Amino acid metabolism pathway metabolites with log 2-fold change greater than 2. **C)** Enriched amino acid biosynthesis and degradation KEGG pathway with respective effect size and NES score across four groups. **D)** Box plot illustrating the log 2-fold changes of all metabolites within the lipid subpathway. **E)** Bar plot illustrating the enrichment of microbial genes in various pathways related to fatty acid biosynthesis and metabolism with respective effect size and NES score. **F)** Microbes with microbial genes with a positive NES score from section C are presented in a plot alongside their corresponding effect sizes across four groups, arranged in sorted order from HG to WU. **I)** Microorganisms carrying microbial genes with a negative NES score from section C are depicted in a chart, displaying their respective effect sizes across four groups, organized in ascending order from HG to WU.

The compounds in amino acids metabolism with the highest abundance (log 2-fold change) in the WU population were: (R)-salsolinol (-18.50), hydroxy-N6,N6,N6-trimethyllysine (-7.63), m-tyramine (-6.43), guanidine (-5.67), 4-imidazoleacetate (-4.86), N2-acetyllysine (-4.51), 3-methylglutaconate (-4.25), N-methylphenylalanine (-4.16) (**Figure 6B, Table S6**). The metabolism of tyrosine, histidine, arginine, proline, lysine, polyamine, methionine, cysteine, taurine, alanine, aspartate, guanidino, and acetamido were higher in the WU population (**Figure 6B, Table S6, S7**). Amino acid pathways enriched in the WU population include methionine biosynthesis, leucine biosynthesis, proline biosynthesis, proline metabolism, tryptophan biosynthesis, shikimate pathway, aromatic amino acid metabolism, isoleucine biosynthesis, histidine biosynthesis, histidine metabolism, arginine and proline metabolism, phenylalanine tyrosine and tryptophan biosynthesis, and valine, leucine and isoleucine biosynthesis (**Figure 6C**). Metabolites of lipids pathways were increased in abundance in the WU population, including androgenic steroids, corticosteroids, endocannabinoid, fatty acid dicarboxylate, fatty acid dihydroxy, fatty acid metabolism acyl, fatty acid monohydroxy, galactosyl glycerolipids, long chain fatty acid, lysophospholipid, lysoplasmalogen, medium chain fatty acid, monoacylglycerol, pregnenolone steroids, primary bile acid metabolism, secondary bile acid metabolism, and sterol (**Figure 6D, Table S8**). The WU population exhibits an abundance of various lipid metabolites, including deoxycholic acid 12-sulfate, chenodeoxycholic acid sulfate, 3-hydroxy sebacate, 1-palmitoyl glycerol (16:0), androsterone sulfate, pregnanediol sulfate, pregnenolone sulfate, 5alpha-androstan-3beta, 17beta-diol mono sulfate, pregnenolone sulfate, and stigmasterol. These lipid metabolites are prevalent in patients with ketoacidosis and are formed from fatty acids through a combination of omega-oxidation, abnormal fatty acid oxidation, and incomplete beta-oxidation. Additionally, they are present in small quantities in commercial food products, plant sterol additions to food, and various components of olive oil and other vegetable oils. Several of these metabolites are natural steroids, belonging to sulfated steroids, neurosteroid, mono-sulfate, and disulfate fractions^77–84^ (**Table S8**). The transcriptomic analysis also showed high fatty acid metabolism and biosynthesis expression, mainly in the WU populations (**Figure 6E**). The WU populations were strongly associated with bacteria involved in the amino acid biosynthesis/metabolism and lipid metabolism pathway (**Figure 6G**).

### Highly diverse carbohydrate metabolism and short-chain fatty acid production in Ju/’hoansi gut microbiome flora

In the Ju/’hoansi gut microbiome, an extensive enrichment of microbial genes associated with various carbohydrate metabolism pathways has been identified (**Figure 7A**). The Ju/’hoansi gut microbiome is highly enriched with microbial genes responsible for carbon fixation, pyruvate oxidation, central carbohydrate metabolism, citrate cycle, the De Ley-Doudoroff pathway, gluconeogenesis, the Embden-Meyerhof pathway, incomplete reductive citrate cycle, the Wood-Ljungdahl pathway, the Arnon-Buchanan cycle, the Calvin cycle, the semi-phosphorylative Entner-Doudoroff pathway, beta-Oxidation, short-chain fatty acids metabolism, and reductive, oxidative and non-oxidative pentose phosphate pathways (**Figure 7A)**. The analysis of carbohydrate metabolites identified several compounds that were either involved in or produced during complex carbohydrate metabolism. Among these metabolites are glucose 6-phosphate, N-acetylmuramate, ribose, N-acetylglucosamine, xylose, mannose, N-acetylneuraminate, ribulose/xylulose, and arabinose, which were found in higher abundance in the rural populations (**Figure 7C**). Rural populations have abundant energy metabolites, including Malate, Fumarate, and Alpha-ketoglutarate (**Figure 7D**). The microbial genes found in the energy generation and carbohydrate metabolism pathways **(Figure 7A)**, prevalent among the Ju/’hoansi, consist of either novel species or species with low to negligible presence in WU populations. Out of 70 abundant species in HG Ju/’hoansi, the top five most abundant ones are: *g_Faecousia;s, CAG-390 sp900753295, UBA1259 sp900760875, UMGS1225 sp900549725, g_Anaerovibrio;s_*(**Figure 7E, Table S9**). The abundant taxa at higher taxonomic levels were the genera Faecousia and Limivicinus. The abundant family comprises Oscillospiraceae, CAG-272, and Ruminococcaceae. At the order level, Oscillospirales, Christensenellales, and Clostridia were the most abundant, with Clostridia being the abundant class.

**Figure 7.**
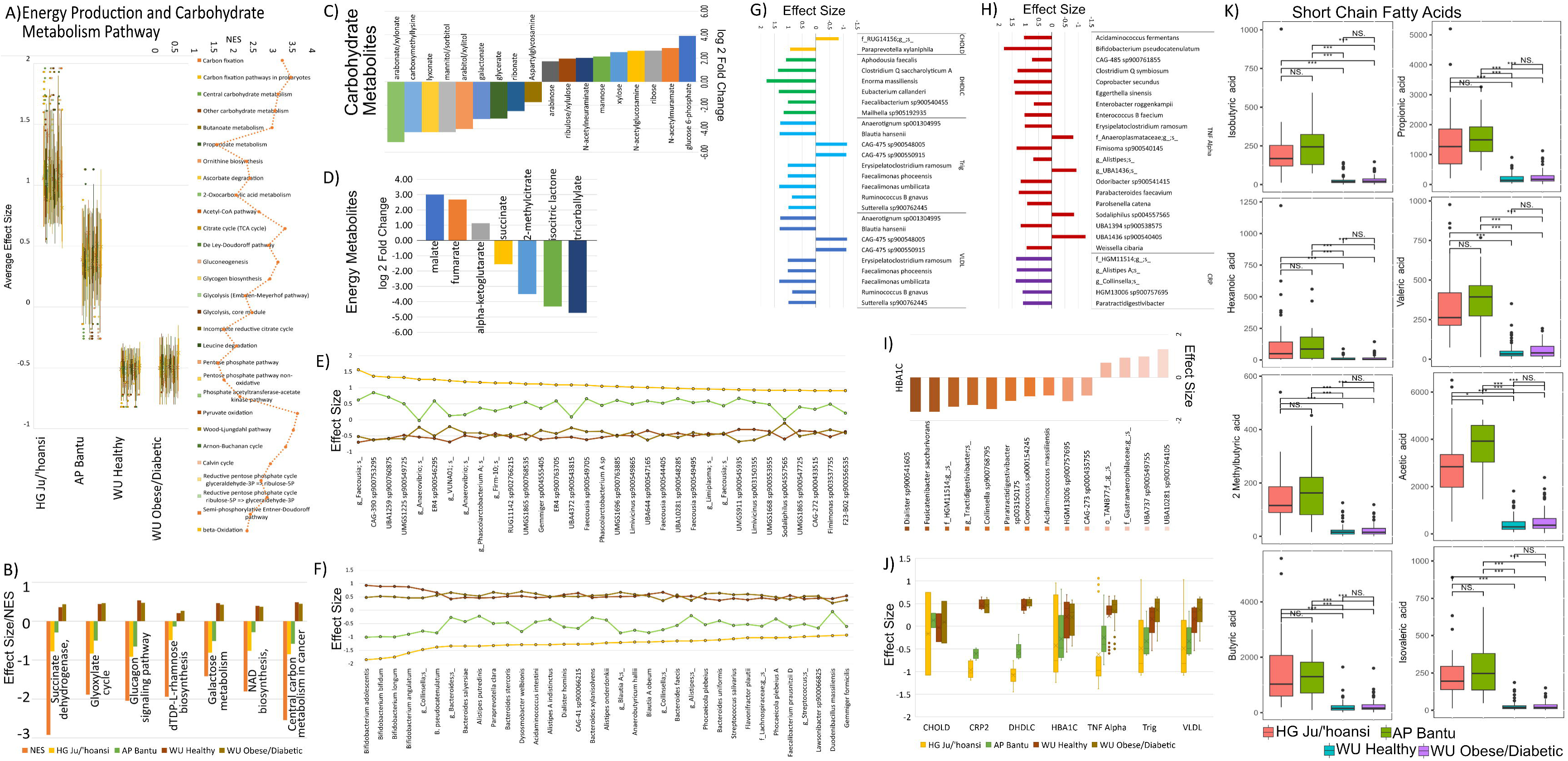
Energy Production Pathway and Blood Biomarker Association: **A)** Multiple energy production pathways highly enriched in the Ju/’hoansi population. **B)** Energy generation pathway enriched in the urban population. **C)** Carbohydrate metabolites and **D)** Energy pathway metabolites abundant in rural and WU populations. **E)** The line plot depicts the effect size of microorganisms carrying microbial genes from section A across four cohorts, sorted based on their abundance in the Ju/’hoansi population. **F)** The chart portrays microorganisms carrying microbial genes falling under section B, showcasing their varying effect sizes within four cohorts. **G)** Microbes associated with lipid profile and **H)** Microbes associated with blood inflammation markers including TNF Alpha and C-Reactive Protein, and **I)** Microbes associated with HbA1c and effect size represent which group of the respective microbe is significantly more abundant in rural or WU. **J)** Box plot of respective microbes’ effect sizes for four cohorts. **E)** Absolute abundance of eight short-chain fatty acids in the four cohorts. The Y-axis represents abundance in terms of µg/g.

The pathways and enzymes enriched in the WU population include succinate dehydrogenase, central carbon metabolism in cancer, glyoxylate cycle, glucagon signaling, dTDP-L-rhamnose biosynthesis, galactose metabolism, and NAD biosynthesis (**Figure 7B**). Multiple metabolites, such as glycerate, galactonate, arabitol/xylitol, mannitol/sorbitol, lyxonate, N6-carboxymethyl lysine, arabonate/xylonate, play a role in advanced glycation end product formation, representing an intermediate step in sugar acid metabolism. These metabolites were notably prevalent in thermally processed foods, sugar alcohols used as sweeteners, milk sugars, and their derivatives, particularly in the WU population ^85–87^ (**Figure 7C**). WU populations had abundant succinate, 2-methylcitrate, isocitric lactone, and tricarballylate energy metabolites (**Figure 7D**). The pathways and enzymes enriched in the WU population are distributed across 48 species **(Figure 7B, Table S9)**. The most abundant species in the WU population are *Bacteroides salyersiae*, *Alistipes putredinis, Acidaminococcus intestini, Bacteroides stercoris, Paraprevotella clara, Alistipes putredinis,* and belong to Bifidobacterium, Collinsella genera (**Figure 7F**).

The data analysis reveals a significant abundance of all seven short-chain fatty acids in rural populations, with a calculated p-value of less than 0.05. Hexanoic acid is especially noteworthy, displaying the highest fold-change at 10.8, indicating a substantial increase. Its mean abundance is reported at 124.29 µg/g in the HG Ju/’hoansi population, 136.52 µg/g in the AP Bantu population, 10.38 µg/g in the WU Healthy group, and 13.66 µg/g in the WU Obese/Diabetic cohort. Isovaleric acid closely follows this trend, exhibiting a fold-change of 8.3 and a mean abundance of 226.72 µg/g in the HG Ju/’hoansi group, 283.59 µg/g in the AP Bantu population, 27.30 µg/g in the WU Healthy cohort, and 30.49 µg/g in the WU Obese/Diabetic cohort. Isobutyric acid also shows a significant fold-change of 7.3, with a mean abundance of 196.70 µg/g in the HG Ju/’hoansi group, 258.99 µg/g in the AP Bantu population, 27.42 µg/g in the WU Healthy group, and 30.93 µg/g in the WU Obese/Diabetic cohort. Valeric acid (reported abundances: HG-324.68, AP-379.64, WU-46.45/56.53) and 2-methylbutyric acid (HG-141.50, AP-177.50, WU-21.35/24.17) demonstrate similar fold-changes at 6.6. Additionally, the three most abundant SCFAs, propionic acid (abundances: HG-1407.98, AP-1566.00, WU-203.14/263.46), acetic acid (abundances: HG-2790.42, AP-3611.25, WU-416.50/564.58), and butyric acid (abundances: HG-1436.47, AP-1309.79, WU-206.48/260.37); all show substantial levels in the rural populations **(Figure 7K)**.

### Ju/’hoansi shows a low abundance of microbes associated with biomarkers and pathways of chronic inflammatory diseases

We utilized Maaslin2 to identify potential microorganisms and pathways that are associated with lipid metabolism, HbA1c, and inflammation. There are 27 microbes that are associated with lipid metabolism, including cholesterol to HDL (high-density lipoprotein) ratio (CHOLD), direct high-density lipoprotein cholesterol (DHDLC), triglycerides, and very low-density lipoprotein cholesterol (VLDL) as shown in **Figure 7G**. Of the 27 identified microbes, only 5 exhibit higher prevalence among the rural population, all distinct microbial strains. Concerning inflammation markers, 20 distinct microbes were found to be linked with TNF Alpha, while 6 microbes were associated with C-reactive protein (CRP). Among the microbes associated with TNF Alpha, only 4 distinct species, 3 of which exhibited an effect size of less than 1.0, were prevalent in the rural population. In contrast, all CRP-associated microbes with effect sizes greater than 1.0 were prevalent in the WU population (**Figure 7H**). Among the 14 microbes associated with HbA1C, only 4 distinct microbes, with an average effect size of 0.9, were prevalent in the rural population despite a high prevalence of carbohydrate-metabolizing microbes in this population. The remaining 10 microbes, with an average effect size of -1.2, were abundant in the WU population (**Figure 7I**). A comparative analysis of effect sizes, depicted in a box plot, reveals that the mean effect size is notably lower in the Ju/’hoansi cohort, followed by the Bantu, WU healthy, and obese/diabetic subjects, in descending order. Significant differences were evident in the abundance of microbes associated with CRP, TNF Alpha, and DHDLC **(Figure 7J)**. These findings imply that most microbes linked to inflammation, lipid, and sugar metabolism were more abundant within the WU population than the rural population.

In order to minimize the possibility of false negatives, we followed a rigorous approach in which only those microbial genes exhibiting statistically significant enrichment in the gene set enrichment analysis (GSEA) pathway undergo thorough testing for their potential association with inflammation markers. By focusing specifically on these enriched genes, we aimed to capture a comprehensive understanding of the relationship between microbial genetic factors and inflammatory processes. The genes we have observed to be enriched in galactose metabolism, glucagon signaling, and central carbon metabolism pathways in cancer have been identified to be associated with TNF Alpha and have significantly higher occurrence among the WU population (**Table S10**). Among the microbial genes associated with CRP abundance, only one gene, the bifunctional 4-hydroxy-3-methylbut-2-enyl diphosphate reductase, present in the isoprenoid and terpenoid biosynthesis pathway, exhibits heightened expression in the rural population. (**Table S10**). Genes that were highly expressed in the WU population were enriched for the central carbon metabolism in cancer, multidrug/antimicrobial resistance, aromatic amino acid metabolism, glyoxylate cycle, lipopolysaccharide metabolism and biosynthesis, cofactor, and vitamin metabolism, pantothenate biosynthesis and shikimate pathways which links metabolism of carbohydrates to biosynthesis of aromatic compounds (**Table S10**). The expression levels of CRP-associated microbial genes were notably lower in the HG Ju/’hoansi population, with subsequent increased expression observed in the AP Bantu and WU populations.

## Discussion

The Ju/’hoansi are among the few remaining groups of hunter-gatherers in the world. While some of their subgroups have transitioned away from a traditional hunter-gatherer lifestyle, their dietary patterns strongly resemble those of their ancestors. This study provides a multi-omics characterization of the gut microbiome of the Ju/’hoansi and compares it to geographically close AP and African descendants from the island of Trinidad in the Caribbean. The findings of this study offer a valuable representation of ancient microbiomes lost due to the impact of Western lifestyles, medicine, and dietary patterns. In addition, this study provides the scientific community with a vast amount of multi-omic data, including metagenomic sequencing reads, metatranscriptomics sequencing reads, global metabolomics data, and detailed metadata to enable secondary analysis by other groups. This research holds significant relevance in elucidating the composition of the primitive microbiome and the impact of industrialized food on the optimal functionality of the gut microbiome.

Our research findings indicate that the Ju/’hoansi population has a significantly greater abundance and diversity of gut microorganisms compared to the geographically proximate AP group, and this distinction is even more pronounced when contrasted with individuals from the Caribbean. Higher bacterial and fungal diversity via 16S and ITS sequencing was also observed in a recent Ju/’hoansi study in the Nyae Nyae Conservancy in northeastern Namibia^25^. The study highlights a significant presence of diverse genera in the HG gut microbiota, demonstrating their crucial role in breaking down complex carbohydrates from plants and producing essential short-chain fatty acids. Notably, several predominant species inhabiting the HG are actively engaged in the metabolism of complex carbohydrates and the generation of energy through various metabolic pathways, encompassing carbon fixation, pyruvate oxidation, the pentose phosphate pathway, the citrate cycle, glycolysis, as well as central carbohydrate metabolic pathways and genes associated with the production of short-chain fatty acids. The high functionality of the HG gastrointestinal microbiome is punctuated by the abundance of specific metabolites such as urolithins, equols, 4-hydroxycinnamate, and feruloylputrescine. These metabolites have been demonstrated to enhance mitochondrial activity and mitophagy, regulate estrogenic activity, and exhibit anti-melanogenic, anti-inflammatory, and antioxidant properties, thereby mitigating the systemic effects of aging and age-related chronic conditions^42–48,88,89^. Numerous beneficial metabolites from bacterial-fungal subpathways were abundant among the Ju/’hoansi. One particular metabolite, 4-acetamidobutanoic acid, derived from the gamma-amino acid pathway, is recognized for its antioxidant and antibacterial properties and its capability to impede the proliferation of pathogens^90^. Our comprehensive multi-omic analysis revealed elevated gene expression of innate prokaryotic defense systems in both the Ju/’hoansi and AP groups, contrasting with increased gene expression of antibiotic-resistant genes in the western groups. This discrepancy may be attributed to the extensive use of antibiotics in the Western diet ^91^. These findings collectively suggest that the Ju/’hoansi population possesses a more intricate metabolic repertoire, potentially contributing to their lower incidence of acute and chronic diseases.

The ingestion of animal products and saturated fats provokes significant alterations in androgen release within the body^92^. On the other hand, a dietary plan that includes higher dietary fiber, reduced intake of refined carbohydrates, decreased consumption of trans and saturated fats, and the addition of omega-3 and omega-9 fatty acids has demonstrated the potential to improve the normalization of the androgen profile. ^93^. Steroid hormones, such as corticosteroids and lipid-based neurotransmitters, including endocannabinoids, play pivotal roles in modulating stress, immune responses, and metabolism. These compounds are integral in regulating energy balance and food intake functions^94,95^. The Ju/’hoansi and the AP groups benefit from a fiber-rich diet while minimizing intake of refined carbohydrates, trans fats, and saturated fats. The metabolic pathways observed in the Trinidad samples predominantly contribute to producing medium- and long-chain fatty acids. Research indicates that these fatty acids can instigate alterations in the composition of gut microbiota, consequently influencing the physiological and immunological aspects of the host^96,97^. The gut microbiota controls the synthesis of these compounds within the intestinal tract. They have the potential to disrupt cellular membranes, induce lipid peroxidation, facilitate the progression of cancer, and modify cellular signaling pathways^98,99^. Many of these compounds are found in vegetable oils, animal fats, and dairy products and are commonly included in small quantities in various commercial Western food items.

In addition to metabolic complexity, the microbiomes of the Ju/’hoansi and AP groups exhibit a significantly increased production of short-chain fatty acids (SCFAs), compared to the obese and healthy cohorts in Trinidad. This heightened SCFA production is attributed to the enzymatic breakdown of carbohydrates by bacteria into simple sugars, which are then fermented to generate SCFAs. This research underscores the potential health implications and benefits associated with understanding and harnessing the metabolic activities of the microbiome. These fatty acids account for up to 10% of the caloric requirements of human cells and play a critical role in maintaining intestinal health. SCFAs are associated with improved gut barrier integrity and enhanced glucose and lipid metabolism. Furthermore, they regulate the immune system, inflammatory response, and blood pressure. In addition, SCFAs play a crucial role in providing energy to colonic cells, promoting the development of tight junctions, facilitating gut regeneration, and exerting direct or indirect influence on gut-brain communication and brain function. Finally, SCFAs enhance T_reg_ activity, improve mitochondrial metabolism, promote keratinocyte differentiation, reduce inflammatory factor expression, and inhibit the inflammatory response^100–103^. The data from our study collectively provides a direct mechanistic link to the lack of reported age-associated chronic inflammatory diseases in the Ju/’hoansi and the AP groups. This suggests a potential avenue for further in-depth exploration of the specific underlying biological mechanisms contributing to this phenomenon in these populations.

The microbiome of the Ju/’hoansi people exhibits a deficiency in microbes associated with fatty acid metabolism, as well as blood inflammation markers TNF Alpha and CRP. The enriched pathway associated with the inflammation marker TNF Alpha, glucagon, regulates glycemia, amino acid, and lipid metabolism, suggesting a higher abundance in urban populations. The CRP-associated enriched 81 pathway, another blood inflammation marker, encompasses crucial elements such as multidrug/antimicrobial resistance, amino acid metabolism, and lipopolysaccharide metabolism/biosynthesis. While the association coefficient between microbes and microbial genes may be small, the results of differential abundance strongly reinforce the connection to lipid metabolism and blood inflammation biomarkers. The study is subject to certain limitations, primarily stemming from the small sample size of the agropastoral population. Despite this, notable differences were observed between the agropastoral and the other three groups. Analysis indicates that the microbiome of the agropastoral population falls between the Ju/’hoansi and the urban population. The observations indicate a potential progression in the microbiome, possibly originating from the rural Ju/’hoansi population and exhibiting an increase or commencing from the urban population and displaying a decrease as it approaches the Ju/’hoansi. Additionally, a limitation of the study involves merging two different metabolomics processed batches, potentially introducing biased analysis. However, the complementary nature of our metatranscriptomics and metabolomics analyses suggests an absence of biased analysis due to the use of different metabolomics batches. It is important to note that the findings from this study are preliminary and would necessitate validation in a case-controlled study using a model organisms or with additional sampling, however, the latter may be restricted by lifestyle, geography and environmental changes.

In summary, our study results provide compelling evidence supporting the unique functionality of the Ju/’hoansi microbiome, which appears to have diminished in the context of industrialized societies. Specifically, we observed distinct microbial profiles associated with the Ju/’hoansi and, to some extent, the AP group that hold potential as live therapeutics for addressing chronic inflammatory diseases prevalent in Western lifestyles. This suggests a potential avenue for targeted interventions. Moreover, this study shows that the differences between healthy and obese/diabetic populations are minimal when compared to the Bantu and the Ju/’hoansi, suggesting that there is a microbial bottleneck for the western microbiome. Our findings also offer valuable insights into the methodologies and datasets necessary for future studies and secondary analyses. These would play a pivotal role in fully elucidating the complex relationships between the microbiome, dietary habits, and environmental factors in the context of health and disease.

### Indigenous community engagement

In order to ensure that scientific discoveries and understanding truly represent all populations rather than being limited to industrialized nations, it is imperative to involve indigenous communities in research endeavors. This approach is crucial to address potential biases and ensure that the benefits of scientific progress are shared equitably among diverse communities, thereby fostering a more inclusive and comprehensive representation of research. In order to maintain ethical standards and prevent exploitation, it is crucial to consider specific considerations when conducting this research. In this study, we performed deep metagenomics, metatranscriptomics sequencing, and metabolomics on anonymized fecal samples collected from Ju/’hoansi and Bantu agropastoralist populations from Namibia in 2019. Dr. Hayes has and continues to have, a close working relationship with the communities, with this study forming a part of a larger volume of work across the region. Specifically, Dr. Hayes has been working with the Ju/’hoansi community participants of this study for over a decade, beginning in 2008. The research presented in the outline of this project has also been communicated back to the community during engagement visits post-COVID-19. Born and educated in neighboring South Africa with a Namibian family, Dr. Hayes is not only well acquainted with the country and cultures, but she also shares a common language with the community, allowing for direct discussions and ensuring a deep level of community involvement and understanding. Dr. Hayes is further supported by a well-established local team, which includes key community members who facilitate site visits and ensure further study translation into the local click-language. As per Ju/’hoansi culture, decisions regarding participation are made at a community rather than at an individual level. The study was conducted without any biases that could have impacted its findings. All eligible participants residing within the focused communities who expressed willingness to participate were included in the study.

## Declarations

### Author Contributions

Managed study participant’s consent and sample collection: VH, RH, HF, JF, DR. Performed computational analysis: HS, JE. Wrote the paper: HS, NG-J. Provided input to the manuscript: VH, SV, KEN. Laboratory work: RR, CK, AA. Conceived, designed, and supervised: HS, NG-J.

### Ethics approval and consent to participate

This study was reviewed and approved by the in-house Institutional Review Board at Garvan Institute of Medical Research (protocol number #280/2017), by the Campus Ethics Committee at the University of the West Indies at St Augustine (reference number: CEC262/08/16) and JCVI (protocol number 2019 277).

### Consent for publication

Informed consent (and assent, where appropriate) and authorization to use, create, and disclose health information for research was obtained from and documented for each research participant enrolled to study at the Garvan Institute of Medical Research and the University of the West Indies at St Augustine, Trinidad.

## Acknowledgments

We in dept to the study participants and their representative communities for their contribution and continued engagement. We are grateful to C.P. Bennett from Evolving Picture in Sydney (https://evolvingpicture.com/) for providing the community recording required to meet local ethical requirements and facilitating with community collections in Namibia, while we acknowledge the community members who have provided further logistical assistance and cultural insights, including B. G/aq’o, N. /kun, J. /kunta, A. Oosthuysen and E. Oosthuysen. We thank the hard working team members of the Department of PreClinical Sciences, University of the West Indies, for their dedicated support.

## Materials and Methods

### Description of sample collection and categorization into groups

In a comprehensive study, we carried out a thorough investigation that involved enrolling and meticulously collecting stool samples from a specific cohort that comprised 61 Ju/’hoansi individuals, who are indigenous people residing in the Kalahari Desert of Namibia, located in southern Africa, as well as 16 agropastoral Bantu individuals living in the immediate vicinity. The collection of samples was undertaken in two phases, with 16 samples obtained in February 2019 and an additional 45 samples gathered in September 2019. These samples were collected for the purpose of the research. Out of the 61 samples, 33 were obtained from female participants, while 28 were obtained from male participants. Additionally, the participants were further categorized based on age, revealing that 30 individuals were over 45 years old and 31 individuals were under 45 years old. The stool samples were collected on-site with meticulous attention to detail and subsequently preserved in liquid nitrogen tanks to ensure their integrity for subsequent analysis. The study protocol received ethical approval from the Garvan Institute of Medical Research in New South Wales, Australia, as well as from JCVI/NIH (2019-277). Dr. Hayes and Dr. Förtsch were granted the necessary permit to collect the samples from the Ministry of Healthy and Social Services (MoHSS Reference: 17/3/3 HEAF).

In Trinidad, a cross-sectional convenience sampling approach was used to recruit study participants characterized as individuals of African-descent (as defined by having at least 3 grandparents of pure African ancestry; *Miller et al. 1989*) who, for the obese/diabetic population, were diagnosed with T2D and/or being obese (BMI≥30kg.m^2;^ n=75); and for the control healthy population (n=75). The study protocol was approved prior to start of recruitment by the following research ethics boards: The University of the West Indies; North Central Regional Health Authority (NCRHA); Southwest Regional Health Authority (SWRHA) and JCVI/NIH. The protocol was modified to adhere to public health guidelines during the Covid-19 pandemic. <u>Inclusion criteria</u>: Individuals, aged 21-60 years of age during consent process, of African-descent (as defined by having at least 3 grandparents of pure African ancestry; *Miller et al. 1989*) who have been diagnosed with type 2 diabetes (T2D) and/or with a BMI≥30kg.m2; and healthy controls.

#### Exclusion criteria

Pregnant women. Individuals with recent antibiotic use; recent history (past 3 months) of gastrointestinal infection or diarrhea; individuals consuming large doses of probiotics. History of: cancer, neurological complications; significant weight loss (>5kg); major dietary changes; chronic alcohol consumption; recent major surgery or hospitalization for more than 24hrs. Positive for Covid-19, HIV, HBV or HCV. History of immunosuppression. Unwillingness to provide blood and stool sample. For controls: in addition to the above: diabetes, markers of metabolic syndrome.

### Recruitment

This was done on a phased basis whereby Controls were recruited first, whilst administrative approvals for visiting health centers were sought to recruit diabetic participants. This phase began during the COVID-19 pandemic, so an electronic flyer detailing the study, its benefits, criteria, and stipend was generated and distributed through the University of the West Indies (UWI) community via its weekly Marketing and Communication electronic newsletter. This flyer was also shared by project members through their social media. Subsequently in the second phase of recruitment a separate electronic flyer was designed specifically to recruit diabetic participants. This second flyer was also circulated by UWI, as above, and in collaboration with the Diabetes Association of Trinidad and Tobago (DATT), shared with its members utilizing the DATT Facebook page. Physical copies of the flyers designed for recruiting participants were placed at high visibility and high traffic areas throughout the UWI as well as in select areas of the North Central Regional Health Authority (NCRHA). During this period nondiabetic controls continued to be recruited through use of the electronic flyers. During Phase 3, we were able to conduct recruitment of diabetic patients through RHA databases. Further, with the end of the global COVID-19 pandemic, restrictions in public health facilities were eased and distribution of physical flyers for diabetic participants was then expanded to include the hospitals and clinics of both the North Central Regional Health Authority (NCRHA) and South West Regional Health Authority (SWRHA) catchments. These physical flyers were also distributed through the private practices of a select few medical practitioners that were willing to do so. The diabetic participants in this study were mainly recruited from health centers and clinics in the NCRHA.

### Screening

Once potential participants agreed to study conditions, having been self-identified or identified by the research group as meeting the study requirements, screening was done to determine eligibility via a project specific screening questionnaire (Appendix A) which was administered either in person or over the phone. If eligibility criteria were met and participants agreed to take part in the study, they were scheduled to make their first visit to the research study site. Screening interviews during recruitment were conducted using standard social distancing protocol (recruiters wore PPE and maintained a distance of 2m). Interview questionnaires also ascertained whether potential participants had, or did have, COVID-19 or flu-like symptoms and/or came into contact with someone who had the disease, and required a history of travel to COVID-19 dense regions in the period between January 2020 and the date of interview.

### Study visit

Once successfully screened, participants were invited. Upon arrival introductions were shared and formally explained the project to the participants. Initial questions were answered. Signed informed consent forms were obtained and stool collection kits were distributed along with the instructions on its use and handling. These kits were red insulated lunch bags, containing two Omnimet gut stool collection kits (DNA Genotek Inc. Ref #ME-200), and a reusable ice pack. Participants were asked to keep the sample cool/frozen once collected. The kits contained a collection swab/spatula, collection vials with stabilizers, and collection assistance paper. After an explanation of the kits and their use, participants were offered a chance to withdraw from the study should this be too difficult or uncomfortable for them.

For sample collection, the participants had an overnight (14 hours) fast then the following morning handed over the collected stool samples. At the study site a questionnaire was administered (Appendix C), along with biometric data collection and blood sampling by the project’s phlebotomist. The participants were initially asked when their last meal was, to ensure they were in a fasting state. After which, the stool samples were checked to ensure both kits were used and were in good condition. Once these checks were done, the study questionnaire was administered to collect demographic information (e.g. age, gender, and occupation). Additionally, baseline medical and medication history were collected such as family medical history, obstetric history, use of medication/supplements and dietary habits. Next, a quick physical examination was completed and included a rapid surface examination and mucous membranes. Participant’s height, weight, blood pressure and heart rate were then recorded using digital and analog scales.

Height was measured using a stadiometer. Blood pressure and heart rate were recorded using an Omron Automatic Blood Pressure Monitor.

### Statistical analyses of sample collection

Data entry: participants were assigned unique identifiers and identifiers were removed from the data files. Participants were assigned to one of two study groups: Obese/Diabetic defined as any participant with BMI ≥30 kg.m^2^, or with an HbA1c value of ≥6.5%; Healthy Controls (denoted as just “Healthy in the manuscript”) were defined as participants with BMI <30kg.m^2^ and not previously diagnosed as type 2 diabetic. All data were validated by a second team member using printouts of the data sheets. Biochemical variables were tested for normality prior to analyses, following which cross tabulation and independent sample t-test were performed to assess significant differences between the two study groups. An adjusted p. value (FDR) < 0.05 were deemed significant. A total of 116 healthy controls responded to advertisements and expressed interest in being a study participant. Of those, 103 responded to follow up and were screened; 78 were deemed eligible and 75 completed the study. A total of 43 obese or type 2 diabetics responded to the advertisements and social media posts; 35 who expressed interest were screened and 27 of those were deemed eligible to participate in the study. The remaining participants in this group were recruited through patient NCRHA data base of patients at the chronic disease clinics in close proximity to the UWI.

### Workstation Decontamination for Isolation and Library Preparation

DNA/RNA isolation and library preparations, the procedures were carried out in a disinfected Class II Biosafety cabinet or Purifier Vertical Clean Bench (PCR Hood) treated with Eliminase (Cat# 1101, Decon Labs Inc., PA, USA) and 70% ethanol afterward exposed to UV light for 30 minutes.

### DNA Extraction of Human Stool Samples

DNA was extracted according to protocol using QIAGEN DNeasy PowerSoil Pro Kit (Cat# 47016, QIAGEN, Hilden, Germany) with mechanically lysed using QIAGEN PowerLyzer 24 Homogenizer (110/220 V) (Cat# 13155, QIAGEN, Hilden, Germany) at 3000 rpm for 30 seconds. After extraction, DNA was eluted in 50uL of C6 buffer (10 mM Tris). Samples were quantified using Qubit HS Assay (Cat# Q32856, ThermoFisher Scientific, Waltham, MA, USA) and DNA quality was measured using an Agilent Genomic DNA TapeStation (Cat# 5067-5366 and 5067-5365, Agilent Technologies, Santa Clara, CA, USA).

### Metagenomic Library Preparation

Metagenomic libraries were generated using the NEBNext Ultra II FS DNA Library Prep Kit (Cat# E7805L, New England Biolab, Ipswich, MA, USA) with the NEBNext Multiplex Oligos for Illumina (Cat# E6448S, New England Biolab, Ipswich, MA, USA). Library prepping was carried out according to protocol for use with gDNA input <100ng. As instructed, all samples were diluted for an estimated 100ng in 26uL volume for starting reaction. Fragmentation was carried out at 37°C for 15 minutes. For adaptor ligation, the NEBNext Adaptor for Illumina were diluted 10-folds in 10 mM Tris-HCl, pH 7.5-8.0 with 10 mM NaCl (recommended for samples input gDNA <100ng). Samples were cleaned using Beckman Coulter Ampure XP beads (Cat# A63881, Beckman Coulter, Pasadena, CA, USA) at a 0.9X ratio. PCR enrichment was carried out by setting the thermal cycler at the following conditions: 98°C for 30 seconds, 98°C for 10 seconds, 65°C for 75 seconds for 4 cycles, 65°C for 5 minutes. Afterwards, libraries were cleaned once more using Ampure XP beads at a 0.9X ratio. Finally, libraries were quantified using Thermo Fisher Qubit 1X dsDNA HS Assay Kit (Cat# Q33231, ThermoFisher Scientific, Waltham, MA, USA) and average fragment size was estimated using Agilent Bioanalyzer High Sensitivity DNA Analysis (Cat# 5067-4626, Agilent Technologies, Santa Clara, CA, USA). Libraries were manually normalized based on the DNA concentration and average fragment size. The pooled library concentration was estimated by qPCR using KAPA Library Quantification kit for Illumina (Kapa Biosystems; Roche Diagnostics Corporation, Indianapolis, IN, USA). Finally, the library was loaded onto a NovaSeq 6000 S2, 300 cycles (2x150 bp) v1.5 as instructed by the manufacturer (Cat# 20028314, Illumina Inc., La Jolla, USA).

### RNA Extraction of Human Stool Samples

RNA was extracted according to protocol using Zymo Quick-RNA Fecal/Soil Microbe Microprep Kit (Cat# R2040, Zymo Research, Irvine, CA, USA). Samples were lysed by bead beating using Vortex Genie-2 set at max speed for 10 minutes. After extraction, RNA was eluted in 15uL of DNase/RNase-Free Water. Samples were quantified using Thermo Fisher Qubit RNA HS kit 100 assays (Cat# Q32852, ThermoFisher Scientific, Waltham, MA, USA). RNA integrity number (RIN) was measured using Agilent Bioanalyzer RNA 6000 nano kit (Cat# 5067-1511, Agilent Technologies, Santa Clara, CA, USA).

### RNA Library Prepping

RNA libraries were generated using llumina’s Stranded Total RNA Prep, Ligation with Ribo-Zero Plus (Cat# 20040529, Illumina Inc., La Jolla, USA) with IDT for Illumina RNA UD Indexes Set A and B (Cat# 20040553 and 20040554, Illumina Inc., La Jolla, USA). Library prepping was carried out according to protocol with some adjustments to fragmentation (fragmentation time was reduced to 30 seconds) and PCR amplification (set to 16 cycles) [RR1] steps because of samples low-quality RNA (the average RIN value was 2.5). Regarding the samples low-quality RNA, input volume was set to a total of 100ng in 11uL starting volume.

After library prepping, samples were quantified using Thermo Fisher Qubit 1X dsDNA HS Assay Kit (Cat# Q33231, ThermoFisher Scientific, Waltham, MA, USA) and average fragment size was estimated using Agilent Bioanalyzer High Sensitivity DNA Analysis (Cat# 5067-4626, Agilent Technologies, Santa Clara, CA, USA). Libraries were manually normalized based on the DNA concentration and average fragment size. The pooled library concentration was estimated by qPCR using KAPA Library Quantification kit for Illumina (Kapa Biosystems; Roche Diagnostics Corporation, Indianapolis, IN, USA). Finally, the library was loaded onto a NovaSeq 6000 S4, 300 cycles (2X150) v1.5 as instructed by the manufacturer (Cat# 20028312, Illumina Inc., La Jolla, USA).

## Metabolomics using the Metabolon Platform

Metabolon received 61 frozen feces and 147 OMNI-met feces samples. Global metabolic profiles comprise a total of 915 compounds of known identity (named biochemicals) were determined. The processing of the samples and the identification of metabolites were carried out by Metabolon in accordance with the specified procedures.

### Sample Accessioning

Following receipt, samples were inventoried and immediately stored at - 80°C. Each sample received was accessed into the Metabolon LIMS system and assigned by the LIMS a unique identifier that was associated with the original source identifier only. This identifier was used to track all sample handling, tasks, results, etc. The samples (and all derived aliquots) were tracked by the LIMS system. All portions of any sample were automatically assigned their own unique identifiers by the LIMS when a new task was created; the relationship of these samples was also tracked. All samples were maintained at -80°C until processed.

### Sample Preparation

Samples were prepared using the automated MicroLab STAR® system from Hamilton Company. Several recovery standards were added before the extraction process’s first step for QC purposes. To remove protein, dissociate small molecules bound to protein or trapped in the precipitated protein matrix, and to recover chemically diverse metabolites, proteins were precipitated with methanol under vigorous shaking for 2 min (Glen Mills GenoGrinder 2000) followed by centrifugation. The resulting extract was divided into five fractions: two for analysis by two separate reverse phase (RP)/UPLC-MS/MS methods with positive ion mode electrospray ionization (ESI), one for analysis by RP/UPLC-MS/MS with negative ion mode ESI, one for analysis by HILIC/UPLC-MS/MS with negative ion mode ESI, and one sample was reserved for backup. Samples were placed briefly on a TurboVap® (Zymark) to remove the organic solvent. The sample extracts were stored overnight under nitrogen before preparation for analysis.

### QA/QC

Several types of controls were analyzed in concert with the experimental samples: a pooled matrix sample generated by taking a small volume of each experimental sample (or alternatively, use of a pool of well-characterized human plasma) served as a technical replicate throughout the data set; extracted water samples served as process blanks; and a cocktail of QC standards that were carefully chosen not to interfere with the measurement of endogenous compounds were spiked into every analyzed sample, allowed instrument performance monitoring and aided chromatographic alignment. Instrument variability was determined by calculating the median relative standard deviation (RSD) for the standards that were added to each sample prior to injection into the mass spectrometers. Overall process variability was determined by calculating the median RSD for all endogenous metabolites (i.e., non-instrument standards) present in 100% of the pooled matrix samples. Experimental samples were randomized across the platform run with QC samples spaced evenly among the injections.

### Ultrahigh Performance Liquid Chromatography-Tandem Mass Spectroscopy (UPLC-MS/MS)

All methods utilized a Waters ACQUITY ultra-performance liquid chromatography (UPLC) and a Thermo Scientific Q-Exactive high resolution/accurate mass spectrometer interfaced with a heated electrospray ionization (HESI-II) source and Orbitrap mass analyzer operated at 35,000 mass resolution. The sample extract was dried then reconstituted in solvents compatible to each of the four methods. Each reconstitution solvent contained a series of standards at fixed concentrations to ensure injection and chromatographic consistency. One aliquot was analyzed using acidic positive ion conditions, chromatographically optimized for more hydrophilic compounds. In this method, the extract was gradient eluted from a C18 column (Waters UPLC BEH C18-2.1x100 mm, 1.7 µm) using water and methanol, containing 0.05% perfluoropentanoic acid (PFPA) and 0.1% formic acid (FA). Another aliquot was also analyzed using acidic positive ion conditions, however it was chromatographically optimized for more hydrophobic compounds. In this method, the extract was gradient eluted from the same afore mentioned C18 column using methanol, acetonitrile, water, 0.05% PFPA and 0.01% FA and was operated at an overall higher organic content. Another aliquot was analyzed using basic negative ion optimized conditions using a separate dedicated C18 column. The basic extracts were gradient eluted from the column using methanol and water, however with 6.5mM Ammonium Bicarbonate at pH 8. The fourth aliquot was analyzed via negative ionization following elution from a HILIC column (Waters UPLC BEH Amide 2.1x150 mm, 1.7 µm) using a gradient consisting of water and acetonitrile with 10mM Ammonium Formate, pH 10.8. The MS analysis alternated between MS and data-dependent MS^n^ scans using dynamic exclusion. The scan range varied slighted between methods but covered 70-1000 m/z. Raw data files are archived and extracted as described below.

### Bioinformatics

The informatics system consists of four major components: the Laboratory Information Management System (LIMS), the data extraction and peak-identification software, data processing tools for QC and compound identification, and a collection of information interpretation and visualization tools for use by data analysts. The hardware and software foundations for these informatics components were the LAN backbone and a database server running Oracle 10.2.0.1 Enterprise Edition.

### LIMS

The purpose of the Metabolon LIMS system was to enable fully auditable laboratory automation through a secure, easy to use, and highly specialized system. The scope of the Metabolon LIMS system encompasses sample accessioning, sample preparation and instrumental analysis and reporting and advanced data analysis. All of the subsequent software systems are grounded in the LIMS data structures. It has been modified to leverage and interface with the in-house information extraction and data visualization systems, as well as third party instrumentation and data analysis software.

### Data Extraction and Compound Identification

Raw data was extracted, peak-identified and QC processed using Metabolon’s hardware and software. These systems are built on a web-service platform utilizing Microsoft’s .NET technologies, which run on high-performance application servers and fiber-channel storage arrays in clusters to provide active failover and load-balancing. Compounds were identified by comparison to library entries of purified standards or recurrent unknown entities. Metabolon maintains a library based on authenticated standards that contains the retention time/index (RI), mass to charge ratio (*m/z)*, and chromatographic data (including MS/MS spectral data) on all molecules present in the library. Furthermore, biochemical identifications are based on three criteria: retention index within a narrow RI window of the proposed identification, accurate mass match to the library +/- 10 ppm, and the MS/MS forward and reverse scores between the experimental data and authentic standards. The MS/MS scores are based on a comparison of the ions present in the experimental spectrum to the ions present in the library spectrum. While there may be similarities between these molecules based on one of these factors, the use of all three data points can be utilized to distinguish and differentiate biochemicals. More than 3300 commercially available purified standard compounds have been acquired and registered into LIMS for analysis on all platforms for determination of their analytical characteristics. Additional mass spectral entries have been created for structurally unnamed biochemicals, which have been identified by virtue of their recurrent nature (both chromatographic and mass spectral). These compounds have the potential to be identified by future acquisition of a matching purified standard or by classical structural analysis.

### Curation

A variety of curation procedures were carried out to ensure that a high quality data set was made available for statistical analysis and data interpretation. The QC and curation processes were designed to ensure accurate and consistent identification of true chemical entities, and to remove those representing system artifacts, mis-assignments, and background noise. Metabolon data analysts use proprietary visualization and interpretation software to confirm the consistency of peak identification among the various samples. Library matches for each compound were checked for each sample and corrected if necessary.

### Metabolite Quantification and Data Normalization

Peaks were quantified using area-under-the-curve. For studies spanning multiple days, a data normalization step was performed to correct variation resulting from instrument inter-day tuning differences. Essentially, each compound was corrected in run-day blocks by registering the medians to equal one (1.00) and normalizing each data point proportionately. In certain instances, biochemical data may have been normalized to an additional factor (e.g., cell counts, total protein as determined by Bradford assay, osmolality, etc.) to account for differences in metabolite levels due to differences in the amount of material present in each sample.

### Batch normalization and Imputation of Data

Normalizing the data is crucial to eliminate instrument batch effects, especially since we have merged data from two different collection samples. This step will ensure the accuracy and integrity of our analysis. For each metabolite, the raw values in the experimental samples are divided by the median of those samples in each instrument batch, giving each batch and, thus, the metabolite a median of one. To remove batch variability, for each metabolite, the values in the experimental samples are divided by the median of those samples in each instrument batch, giving each batch and, thus, the metabolite a median of one. For each metabolite, the minimum value across all batches in the median scaled data is imputed for the missing values. For each sample, the Batch-normalized data is divided by the value of the normalizer. Then, each metabolite is re-scaled to have a median = 1 (divide the new values by the overall median for each metabolite). Then, imputation is performed. The batch-norm-imputed data is transformed using the natural log. Metabolomic data typically displays a log-normal distribution; therefore, the log-transformed data is used for statistical analyses.

### Assembly, classification, and dereplication for metagenome-assembled genomes

We used VEBA v1.2.0 to pursue a metagenomics approach by assembling and binning each metagenome separately, performing species-level clustering, and a within SLC orthology analysis ^26,104^. Using VEBA’s assembly.py module, each metagenome was assembled per sample using SPAdes v3.15.2 (metaSPAdes mode)^105^. Viruses were identified and classified using geNomad v1.3.3 ^106^ (VEBA’s binning-viral.py and classify-viral.py modules). Prokaryotes were binned using several binning algorithms, including: 1) MaxBin2 v2.2.7^106^, 2) Metabat2 v2.15^107^, and 3) CONCOCT^108^. The binning assignments were refined using DAS Tool v1.1.2^109^, yielding 9,011 MAGs that were classified using GTDB-Tk v2.2.3 (reference database: R207^27^) and quality assessed using the lineage_wf pipeline of CheckM v1.1.3^110^ (VEBA’s binning-prokaryotic.py and classify-prokaryotic.py modules). Eukaryotic genomes were binned using Metabat2 v2.15, quality assessed using BUSCO v5.4.3^111^, and classified against VEBA’s microeukaryotic protein database (VEBA’s binning-eukaryotic.py and classify-eukaryotic.py modules). Both prokaryotic and eukaryotic genomes were only used if they were at least medium quality according to the guidelines established by the Genomic Standard Consortium (>50% completeness and <10% contamination;^112^. The recommended cutoffs in CheckV for considering viral contigs are the following: 1) the number of viral genes > 0; 2) the number of viral genes ≥ 5 times the number of host genes; 3) completeness ≥ 50%; 4) checkv_quality and miuvig_quality are above medium quality^113^. Only non-proviruses were used in downstream analysis to prevent host bias.

Genome and protein-level clustering was performed using VEBA’s cluster.py module. The dereplication of redundant species was clustered using FastANI v1.32^114^ with a cutoff of ≥ 95% ANI; as the authors recommended, connected components were determined using NetworkX v2.5. FastANI clustering was separately prokaryotic, eukaryotic, and viral organisms to yield SLCs. For each SLC, the proteins within the pangenome were clustered into SSPCs using MMseqs2 with a query coverage of 80% and percent identity of 50%^115^.

### Gene models, functional annotation, and orthology analysis

VEBA’s binning modules handle all the gene modeling in the backend. To be specific, gene models were created for prokaryotic genomes using Prodigal v2.6.3 (--meta mode;)^116^, viral genomes using Prodigal-GV v2.10.0^106^, and eukaryotic genomes with MetaEuk generating putative proteins and GFF files used for annotations and read mapping^117^. The putative proteins were annotated using both Diamond v2.1.7^118^ against UniRef90^119^ and KofamScan v1.3.0^120^.

### Read preprocessing and mapping to the genome catalog

Using VEBA’s index.py and mapping.py, we aligned reads to the contigs for each genome to assess abundance and expression for metagenomics and metatranscriptomics datasets, respectively. Bowtie2 v2.5.2^121^ was used for all read mapping to MAGs. Read counts tables were constructed with featureCounts v2.0.6^122^ using the bam files generated from Bowtie2, the assembly fasta, and the gene models. Counts for SLC or MAGs were computed by summing contig-level read counts using a SAF file generated from MAGs.

Reads were preprocessed using VEBA’s preproces.py, which is a wrapper around Fastq Preprocessor. In the backed, reads were preprocessed using FastP v0.23.4^123^ to quality trim and remove adapters. Quality trimmed reads were to the human genome (GRCh37.p4) via Bowtie2 v2.5.2 and removed aligned reads for assembly and mapping. Reads were quantified using SeqKit v2.3.1^124^.

### Statistical Analyses of Gut Microbiome Data

Differential (relative) abundance metrics fail to address compositionality. Extensive tool benchmarking has demonstrated the susceptibility of commonly employed differential (relative) abundance tools to sparsity, leading to unacceptably high rates of false positive identifications. It is imperative to consider high-throughput sequencing-derived microbiome study datasets as compositions throughout the analysis process. To circumvent the challenges associated with relative abundance analysis, we conducted all microbiome analyses using Compositional Data Analysis (CoDA) with ALDEx2. The analysis relied on the ALDEx2 (ANOVA-Like Differential Expression) package version 1.34 in R 4.3.2125 to identify differential abundances in the gut microbiota at a species level. For a rigorous effect size calculation in ALDEx2, we obtained analysis using 1,000 Dirichlet Monte-Carlo Instances (DMC) at 0.75 gamma. The identical parameters were employed to identify differentially abundant microbial genes/orthologs. In comparing rural and urban (WU) settings, merging HG and AP datasets results in the formation of rural datasets, while the combination of WU healthy and WU obese/diabetic datasets yields the WU datasets. However, it is important to note that all four datasets were categorized as separate groups to calculate each group’s effect size. The ALDEx2 tool is a powerful solution for conducting statistical tests on the centered log-ratio (clr) values obtained from a modeled probability distribution of the dataset. This tool effectively addresses the issue of false-positive identification by providing expected values of parametric and non-parametric statistical tests, along with effect-size estimates. It has demonstrated a remarkable ability to minimize this problem to near-zero levels in real and modeled microbiome datasets, proving its robustness even when dealing with dataset subsets^125,126^. Among the various methods available for analyzing microbial differential abundance, Aldex2 stands out for its exceptional performance in controlling the false discovery rate (FDR)^127^.

We utilized the gseKEGG function within the clusterProfile v4.10.0 R package to perform the KEGG pathway gene set enrichment analysis^28^. This involved identifying significantly enriched KEGG pathways associated with the differentially abundant microbes. For the enrichment analysis, we used all KEGG orthologs present in the differentially abundant microbes as the background genes, ensuring a comprehensive and informative pathway enrichment analysis. The KEGG pathway, modules, and BRITE information were obtained through the KEGGREST R package, an R interface to the KEGG database^128^. In the context of metabolomics data analysis, differentially abundant metabolites were identified using the t-test with a false discovery rate (FDR) threshold set at less than 0.05. The fold changes are determined by computing the ratios between the rural and WU populations. All of the analyses in the current study were performed at adjusted p.value < 0.05.

